# The legacy of recurrent introgression during the radiation of hares

**DOI:** 10.1101/2020.06.19.160283

**Authors:** Mafalda S. Ferreira, Matthew R. Jones, Colin M. Callahan, Liliana Farelo, Zelalem Tolesa, Franz Suchentrunk, Pierre Boursot, L. Scott Mills, Paulo C. Alves, Jeffrey M. Good, José Melo-Ferreira

## Abstract

Hybridization may often be an important source of adaptive variation, but the extent and long-term impacts of introgression have seldom been evaluated in the phylogenetic context of a radiation. Hares (*Lepus*) represent a widespread mammalian radiation of 32 extant species characterized by striking ecological adaptations and recurrent admixture. To understand the relevance of introgressive hybridization during the diversification of *Lepus*, we analyzed whole exome sequences (61.7 Mb) from 15 species of hares (1- 4 individuals per species), spanning the global distribution of the genus, and two outgroups. We used a coalescent framework to infer species relationships and divergence times, despite extensive genealogical discordance. We found high levels of allele sharing among species and show that this reflects extensive incomplete lineage sorting and temporally layered hybridization. Our results revealed recurrent introgression at all stages along the *Lepus* radiation, including recent gene flow between extant species since the last glacial maximum, but also pervasive ancient introgression occurring since near the origin of the hare lineages. We show that ancient hybridization between northern hemisphere species has resulted in shared variation of potential adaptive relevance to highly seasonal environments, including genes involved in circadian rhythm regulation, pigmentation, and thermoregulation. Our results illustrate how the genetic legacy of ancestral hybridization may persist across a radiation, leaving a long-lasting signature of shared genetic variation that may contribute to adaptation within and among species.

---

Species radiations are often accompanied by extensive gene flow between nascent lineages (e.g., Lamichhaney et al. 2015; Árnason et al. 2018; Malinsky et al. 2018; Li et al. 2019; Barth et al. 2020). Genetic signatures of hybridization between several closely related species could either represent recent or ongoing introgressive hybridization (Eaton et al. 2015), or the remnants of hybridization among ancestral populations that remain shared among contemporary species (Malinsky et al. 2018; Li et al. 2019). Although these alternatives can be difficult to differentiate in large radiations (Eaton et al. 2015; Malinsky et al. 2018; Vanderpool et al. 2020), both ancient and contemporary introgression has been linked to local adaptation in several systems (e.g., Liu et al. 2015; Gittelman et al. 2016; Meier et al. 2017; Barlow et al. 2018; Giska et al. 2019; Svardal et al. 2020). Thus, unraveling the tempo and contribution of introgression to standing genetic variation within and among species remains a critical step in understanding the overall importance of introgression to evolution.

Reconstructing the history of hybridization between several closely related species requires inferring evolutionary relationships among species while considering the two primary processes – incomplete lineage sorting and gene flow – that may cause sharing of genetic variation among populations (Malinsky et al. 2018). The network multispecies coalescent (NMSC) model (Than et al. 2011; Solís-Lemus et al. 2017; Degnan 2018) offers one promising framework that appears to resolve species relationships in the face of multiple reticulation events and rapid speciation (Kozak et al. 2018; Edelman et al. 2019). However, the NMSC is still prohibitive for large datasets and choosing the exact number of migration events is not straightforward (Yu and Nakhleh 2015). Alternatively, site-based summary statistics based on tree asymmetries (e.g., Green et al. 2010; Pease and Hahn 2015), or admixture proportions (e.g., Reich et al. 2009; Martin et al. 2015; Malinsky et al. 2018) are simpler to implement, but offer less power for localizing the timing and number of introgression events when recurrent hybridization is layered across a phylogeny (Malinsky et al. 2018). A combination of methods is thus most appropriate to infer a species tree that may have layered events of hybridization throughout time (e.g., Kozak et al. 2018; Malinsky et al. 2018; Edelman et al. 2019; Li et al. 2019).

Hares and jackrabbits comprise a group of 32 species (genus *Lepus*; Smith et al. 2018) whose common ancestor likely originated in North America and spread throughout most of the Northern Hemisphere and Africa presumably in the last 4-6 million years (Yamada et al. 2002; Matthee et al. 2004; Melo-Ferreira et al. 2012). Hares are primarily associated with open grasslands, but can be found across a broad range of biomes (e.g., desert, forest, or arctic) and elevations (e.g., from sea level to the Himalayan or Ethiopian plateau; Smith et al., 2018). The *Lepus* radiation also provides case studies of hybridization and introgression, since admixture has been detected among several modern species pairs (e.g., Liu et al. 2011; Melo-Ferreira et al. 2012; Tolesa et al. 2017; Jones et al. 2018; Seixas et al. 2018; Lado et al. 2019; Kinoshita et al. 2019). Selection on introgressed variation has been hypothesized to have aided the range expansion of the Iberian hare (Seixas et al. 2018), and has been directly linked to convergent adaptive evolution of non-white winter coats in populations of two species that change the color of their pelage seasonally (Jones et al. 2018; Giska et al. 2019; Jones et al. 2020a). These studies suggest that the relatively recent exchange of genetic variation among extant *Lepus* species has provided an important source of adaptive variation. However, the phylogenetic relationships among *Lepus* species remain poorly resolved (Halanych et al. 1999; Matthee et al. 2004; Melo-Ferreira et al. 2012; Melo-Ferreira and Alves 2018), and the contribution of ancient gene flow to the *Lepus* evolutionary history in a deeper phylogenetic context remains unknown.

Here, we use exome-wide data to infer the evolutionary history of 15 *Lepus* species and show that hybridization between lineages has likely occurred since the origin of the radiation. The combination of incomplete lineage sorting and these temporally layered events of hybridization have resulted in extremely high levels of shared genetic variation among extant species, including species that currently occur on different continents. We then use the case of ancient admixture among northern latitude species that occupy highly seasonal environments to investigate the gene content and possible functional relevance of introgressed genomic regions. Our work demonstrates that recurrent introgression throughout evolutionary history has made a substantial contribution to genetic variation within and among species of this widespread mammalian radiation.

## Materials and Methods

### Taxon Sampling and Exome Sequencing

We generated new genome-wide resequencing data targeting 207,691 exonic and non-coding regions [totaling 61.7 Megabases (Mb) total] from 14 hare species (30 individuals) and the outgroup pygmy rabbit (*Brachylagus idahoensis*; 2 individuals). We combined these data with published whole exomes from four snowshoes hares [*Lepus americanus*; NCBI Sequence Read Archive BioProject PRJNA420081 from Jones et al. (2018, 2020b)] and extracted data from the reference genome of European rabbit (*Oryctolagus cuniculus*; OryCun2.0; Carneiro *et al*., 2014) to use as second outgroup. Our total sample of 15 hare species (34 individuals, 1 to 4 individuals per species) and 2 outgroup species (3 individuals) included species from all major regions of the *Lepus* native distribution: Africa (3 species), Africa and Eurasia (1 species), Eurasia (6 species) and North America (5 species) (Fig. 1 and Supplementary Table S1).

**Fig. 1.**
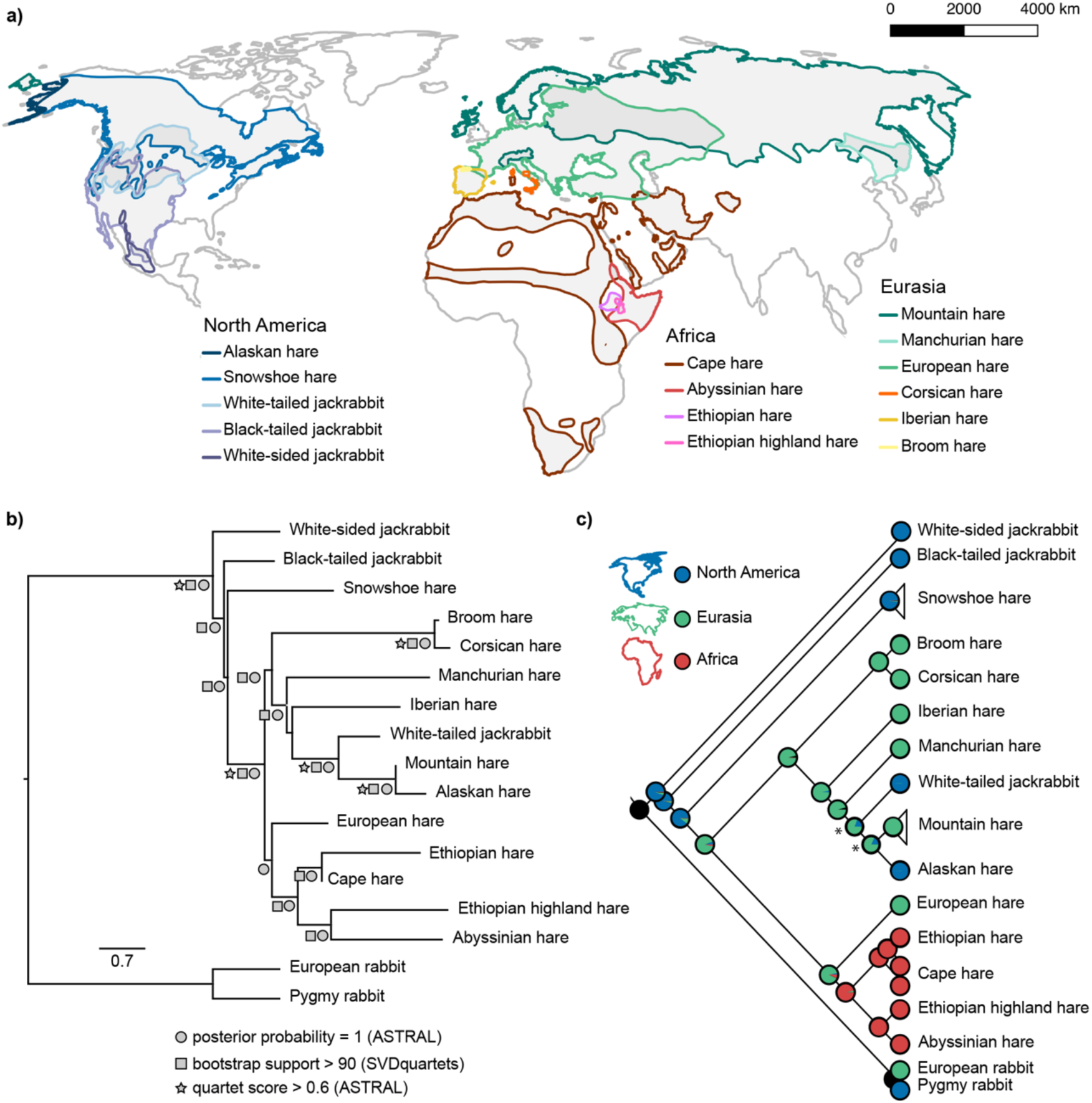
Hare (*Lepus* spp.) evolutionary history and biogeography. **A)** The distribution of the 15 hare species studied here; **B)** Coalescent species tree estimated with *ASTRAL;* nodes with *ASTRAL* posterior probability equal to one, *ASTRAL* quartet score higher than 0.6, and *SVDquartets* bootstrap support higher than 90 are labeled; **C)** Maximum-likelihood phylogeny of *Lepus* showing ancestral range reconstructions. Pie charts show proportional likelihoods that the common ancestral was distributed in North America (blue), Eurasia (green) or Africa (red). Asterisks depict nodes with equivocal reconstruction of ancestor distribution.

Exome capture experiments were performed following the procedures outlined in Jones et al. (2018) and in the Appendix. Briefly, we obtained samples as tissue or extracted DNA including samples from previous studies or through loans from collaborators (Supplementary Table S1). Depending on the sample, genomic DNA was isolated using a saline extraction method (Sambrook et al. 1989) or DNeasy Blood & Tissue Kit (Qiagen) (Supplementary Table S1 and Appendix). We prepared Illumina sequencing libraries for each sample following Meyer and Kircher (2010) with minor modifications [see Appendix and Jones et al. (2018)]. Sequencing libraries were then enriched using NimbleGen SeqCap EZ v.4.3 protocol and a custom capture design consisting of 213,164 probes targeting ∼25 Mb of protein-coding exons, ∼28 Mb of untranslated regions, and ∼9 Mb of intron/intergenic regions (Jones et al. 2018). Hybridization reactions were performed in two separate equimolar pools of indexed libraries (31 and 29 libraries, Supplementary Table S1), together with samples used for other studies. The target enriched pools were each sequenced across two lanes of an Illumina HiSeq1500 sequencer (125 bp paired-end reads) at CIBIO-InBIO’s New-Gen sequencing platform, Portugal.

### Read Processing and Genotyping

We trimmed adapters, low-quality bases, merged overlapping reads, and removed PCR duplicates from raw reads using the *expHTS* pipeline (v.0.Mar112016; https://github.com/msettles/expHTS). We then used *pseudo-it* (v1; Sarver et al. 2017) to generate pseudo-reference exomes for each species by iteratively mapping (four iterations and allowing ambiguity codes) cleaned single and paired-end reads from one individual per species (Supplementary Table S1) to the rabbit reference genome (OryCun2.0) (Carneiro et al. 2014). For snowshoe hares and black-tailed jackrabbits, we used pseudo-references generated by Jones et al. (2018).

We then mapped data from all individuals to each species-specific pseudo-reference using *bwa-mem* (v.0.7.12-r1039; Li 2013) with default options. Mapped reads were sorted with *samtools* (v1.4; Li et al. 2009), assigned to read groups, filtered for duplicates (*Picard* v1.140; http://broadinstitute.github.io/picard/), and realigned for insertion-deletion length variation using GATK (v3.4.46; Van der Auwera et al. 2013). We calculated coverage statistics and capture efficiency using *CalculateHSMetrics* from *Picard*. For each individual, we called and filtered genotypes using the *bcftools* (v1.4; Li 2011) mpileup, call and filter pipeline and used the curated genotypes to construct consensus exome fasta sequences in the OryCun2.0 coordinate system (see Appendix for more details on the filtering pipeline). Many of the steps of the bioinformatics pipeline in our analyses benefited from parallelization provided by GNU parallel (Tange 2011).

### Species Tree Inference

We used a concatenated alignment to estimate a single bifurcating phylogeny using a maximum likelihood (ML) search and rapid bootstrapping run under the GTR+Γ model of sequence evolution (autoMRE option) in *RAxML* (v8.2.10; Stamatakis 2018). We then used two complementary methods to infer multispecies-coalescent trees while accounting for local variation in phylogenetic histories along the genome. First, we extracted alignments from the targeted regions and 200 base pairs (bp) of flanking sequences across 50 kilobase (kb) genomic intervals, considering a balance between the expected extent of LD in hares (10-20 kb; Jones et al. 2018) and the retention of information for phylogenetic analysis (alignment length > 1 kb). For each window, we filtered positions with missing data for > 30% of individuals using *TriSeq* (*TriFusion 1*.*0*.*0; http://odiogosilva.github.io/TriFusion/), excluded windows smaller than 1 kb, and used RAxML* to estimate local maximum likelihood trees (GTR+Γ, 100 bootstraps) and corresponding tree certainty scores. We then used all windows with tree certainty scores above 5. The tree certainty score is the sum of certainty scores for all internodes of a tree. The internode certainty score weighs the support of the bipartition represented by a given internode against the support of the second most prevalent conflicting bipartition (Salichos et al. 2014). In our case, the maximum theoretical value of the tree certainty score is 31, or k – 3 with k equal to the number of taxa (Salichos et al. 2014). We unrooted the gene trees using R package *ape* (Paradis et al. 2004) to estimate a consensus species tree using *ASTRAL-III* (5.6.3; Zhang et al. 2018).

We also estimated coalescent species trees using only variable sites with *SVDquartets* (Chifman and Kubatko 2014) implemented in *PAUP\** (4a163; Swofford 2003). For the analyses based on variable sites, we recovered single nucleotide variants (SNVs) distanced 10 kb along the genome, and excluded sites with missing information for >30% of the individuals. For both *SVDquartets* and *ASTRAL* analyses, species trees were estimated with and without assigning species identities and using sites/intervals genome-wide or only from the X chromosome. The European and/or pygmy rabbits were included for all analyses requiring outgroups. Additional details on the phylogenetic analyses are provided in the Appendix.

### Bayesian Divergence Time Inference

We performed Bayesian inference of divergence times in the inferred species tree using an approximate maximum likelihood method and assuming an autocorrelated relaxed molecular clock, implemented in *MCMCtree* (*PAML v*.*4*.*9*; Yang 2007) and described in dos Reis and Yang (2011, 2019). For one individual per species (Supplementary Table S1), we extracted the coding sequence for all genes included in our capture design (18,798 genes in the OryCun2.0 ENSEMBLE 94 database), selecting the longest transcript per gene using R package *biomaRt* (v2.34.2; Durinck et al. 2005, 2009) and excluding alignments with more than 20% missing data using *AMAS* (Borowiec 2016) (see Appendix for details). With these, we constructed a concatenated alignment with three partitions, corresponding to the three codon positions. We assumed GTR+Γ for the model of sequence evolution, we used the prior of 3.33 for the average substitution rate per site per 100 million years, following Matthee et al. (2004). We discarded the first 1,000,000 samples as burn-in and ran the program until we gathered 1,000,000 samples from the posterior, sampling every 10 iterations, and repeated the analysis twice to ensure convergence. We checked for convergence between the two runs by confirming a linear correlation between posterior times, trendless trace plots, and high effective sample size (ESS) values following dos Reis and Yang (2019). Finally, we also checked for a linear relationship between posterior times and confidence interval widths in infinite sites plots (Inoue et al. 2010).

*Lepus* is poorly represented in the fossil record. The earliest hare record dates to the early Pleistocene (2.5 million years ago; Ma) (White 1991; Lopez-Martinez 2008), which is much more recent than molecular estimates for the genus extrapolated from deeper fossil calibrations (4-6 Ma; Yamada et al. 2002). Therefore, we dated the species tree either (1) using previous molecular estimates of 4-6 Ma for *Lepus* diversification extrapolated from deep fossil record calibrations of the order Lagomorpha (Yamada et al. 2002) and 9.7-14.5 Ma for the *Oryctolagus*-*Lepus* divergence (Matthee et al. 2004) or (2) using the fossil estimates of 2.5 Ma for the lower bound of *Lepus* diversification, and constraining this node to be no older than 4.8 Ma, which is when the fossil record suggests that the common ancestor of Leporids (rabbits and hares) existed (Hibbard 1963; White 1991).

### Ancestral Range Reconstruction

We reconstructed ancestral continental ranges for *Lepus* using the likelihood reconstruction method (Schluter et al. 1997; Pagel 1999) implemented in *Mesquite* (v3.51; Maddison and Maddison 2018). We generated a character matrix coding each species as being distributed in North America (0), Eurasia (1) or Africa (2). Even though the Cape hare *sensu lato* is distributed in Africa and Eurasia, Lado et al. (2019) showed deep divergence and non-monophyly of African and Eurasian lineages. Because our Cape hare samples represent the African lineage, we assigned the Cape hare distribution to Africa. We used the Markov k-state one-parameter model (Mk1) (Lewis 2001) and the rooted topology and branch-lengths of the concatenated ML tree to infer ancestral states of internal nodes. Nodes for which the decision threshold differed by less than 2.0 between states were considered ambiguous.

### Gene Tree Discordance and Phylogenetic Networks

To explore the amount and effect of gene tree discordance in our dataset, we performed a series of analyses. First, we produced a split network using all gene trees previously used as input for *ASTRAL* (tree certainty scores > 5) using *SplitsTree4* (v4.14.6; Huson and Bryant 2006) with the option “Consensus Network with distances as means” and a 5% threshold. We also used the same set of gene trees (tree certainty scores > 5) to produce a majority rule consensus tree with *RAxML* (-L MRE option), including internode certainty scores (Salichos et al. 2014). We then used *DiscoVista* (Sayyari et al. 2018) to plot *ASTRAL* quartet frequencies around nodes of interest. Finally, we calculated Robinson-Foulds normalized distances between gene trees and the *ASTRAL* species tree where individuals are not assigned to species, and among gene trees using the function *RF*.*dist()* from the R package *phangorn* (2.4.0 ;Schliep 2011). This metric varies between 0 (no discordance between trees) and 1 (complete discordance).

We used *PhyloNet* (v3.6.6; Yu and Nakhleh 2015) to model species relationships under the network multispecies coalescent model, using all local genealogies (tree certainty scores > 5). We ran *InferNetwork_MPL* (Yu and Nakhleh 2015) with 0 to 4 migration events, associating individuals to species (option -a), and optimizing branch lengths and inheritance probabilities to compute likelihoods for each proposed network (option -o). We used the best likelihoods per run to calculate Bayesian Information Criteria (BIC) and Akaike information criteria corrected for small samples sizes (AICc) following (Yu et al. 2012, 2014) to compare the resulting networks. Networks were visualized with *IcyTree* (https://icytree.org; last accessed July 2019). We also reconstructed ancestral population graphs with *TreeMix* (v1.13, options -global, -noss and -se; Pickrell and Pritchard 2012) based on the SNV dataset used for *SVDQuartets*. We allowed 0 to 9 migration events and used the white-sided jackrabbit (*Lepus callotis*) as the outgroup relative to all other *Lepus* species (see *Results*).

### Genetic Diversity, Divergence, and Admixture

We used the *genomics general* toolkit (https://github.com/simonhmartin/; last accessed January 14, 2019) to estimate pairwise genetic distances between species and nucleotide diversity within species, and a custom script (available at https://github.com/evochange) to calculate the number of heterozygous sites per individual and the subset shared among species. All diversity estimates were based on a genome-wide concatenated alignment, where we excluded sites with missing information for >30% of the individuals.

We then used *genomics general* and custom scripts (available at https://github.com/evochange) to calculate several variants of the D-statistics (Green et al. 2010) from the informative sites in the same filtered alignment, treating the European rabbit sequence as the ancestral state (additional details are provided in the Appendix). Briefly, we calculated the minimum absolute value of D (D_min_) (Malinsky et al. 2018) for all possible species trios. We calculated z-scores for each D value using a 1 Mb block jackknife approach. After finding the minimum D per trio, D values with Bonferroni-corrected *P* ≤ 0.05 were considered significantly different from zero. We then calculated the ‘f-branch’ statistic (*f*_*b*_*(C)*) (Malinsky et al. 2018). The ‘f-branch’ statistic measures admixture proportion between species *C* and branch *b* by calculating admixture proportion among all possible *f*(A,B,C,O) combinations where *A* are all descendants of branch *a* (sister to *b*), *B* are all descendants of branch *b*, and *C* is the donor taxa. *f*_*b*_*(C)* is the minimum *f* value across all possible *B* and the median across all possible *A*. A significant *f*_*b*_*(C)* value means that all descendants *B* of branch *b* share alleles with *C*, which is more parsimoniously explained by an event of ancestral introgression from *C* to *b* (Malinsky et al. 2018). Using the inferred species tree, we determined all conformations (A,B;C,O) needed to calculate *f*_*b*_*(C)* for all pairs of *C* species and *b* branches following Malinsky et al. (2018), custom scripts (available at https://github.com/evochange) and R package *treeman* (1.1.3; Bennett et al. 2017). For each conformation, we calculated ‘admixture proportion’ (f_G_) as defined in Martin et al. (2015) and Malinsky et al. (2018) and z-scores with 1 Mb block jackknife approach following Malinsky et al. (2018). We also calculated f_hom_ (Martin et al. 2015) between black-tailed jackrabbits (P3) and several snowshoe hare populations (as P1 and P2) to evaluate levels of admixture estimated with *f*_*b*_*(C)* for this species pair (see *Results* and *Discussion*).

We also estimated the fraction of admixture (*f*_*d*_) (Martin et al. 2015) across 50 kb genomic sliding windows (≥100 sites, 5 kb steps), to localize introgression in the genomes of northern latitude species (snowshoe hares, mountain hares, Alaskan hares, and white-tailed jackrabbits). We performed three scans testing introgression between snowshoe hares (P3) and Alaskan hares, mountain hares or white-tailed jackrabbits (alternative P2), using the Iberian hare as P1. Windows of top 0.5% *f*_*d*_ were considered significant. Following Liu et al. (2015), we considered that significant windows in all three tests reflected introgression between snowshoe hares and the ancestral lineage of white-tailed jackrabbits/mountain hares/Alaskan hares, while significant windows in only one test result from recent introgression between the focal extant species. We obtained the annotation of genes in these windows from the rabbit reference and performed an enrichment analysis in *g:Profiler* (accessed September 2019; Raudvere et al. 2019) using default parameters. We also calculated d_xy_ for *f*_*d*_ outlier windows and the exome-wide d_xy_ distribution between the focal pair of P2-P3 species.

## Results

### Whole Exome Sequencing Data

We analyzed whole exome sequence data (61.7Mb) from 15 hare species and two outgroups, combining newly generated (30 individuals from 14 hare species and 2 individuals from the outgroup pygmy rabbit) and published data (4 individuals from 1 additional hare species and the outgroup European rabbit; Carneiro et al. 2014; Jones et al. 2018, 2020b) (Supplementary Table S1). Custom DNA captures showed high efficiency (32.3 average fold-enrichment) and specificity (average 10% of sequenced bases off-target; Supplementary Table S2). Mapping cleaned reads onto species-specific pseudo-references resulted in an average target sequencing coverage of 16× (5-35× on average per sample; Supplementary Table S2) with 57.8 million genotyped sites per individual (Supplementary Table S2). Two lower coverage individuals (one hare individual and one pygmy rabbit individual) and one locality duplicate (one hare) were removed from the final dataset to maximize data quality and avoid geographic redundancy (Supplementary Table S1). All analyses were performed on a dataset of 15 hare species (32 individuals), one pygmy rabbit, and the rabbit reference genome, unless otherwise noted (see Supplementary Table S1).

### Phylogenetic Relationships among Hares

The overall topologies of the concatenated ML phylogeny (11,949,529 positions with no missing data) and the multispecies-coalescent species trees of *ASTRAL* (8,889 gene trees estimated from 50 kb genomic intervals; alignment lengths between 1 kb and 29 kb) and *SVDquartets* (45,779 unlinked SNPs) were largely concordant (Supplementary Figs. S1, S2 and S3). Branching relationships were highly supported in general (*ASTRAL* posterior probabilities equal to 1 and *SVDquartets* bootstrap supports > 90), except for i) the placement of European hares (*L. europaeus*) as sister to the clade of African species, and ii) the sister relationship of the Iberian hare (*L. granatensis*) and the clade containing the white-tailed jackrabbit (*L. townsendii*), Alaskan and mountain hares (Fig. 1 and Supplementary Figs. S1, S2, S3 and S4). Even though the taxonomy of some hare species has been controversial (e.g., Halanych et al. 1999; Alves et al. 2008; Melo-Ferreira and Alves 2018; Lado et al. 2019), our analyses recovered most species as monophyletic (Supplementary Figs. S1, S2a and S3a). Possible exceptions were the potential paraphyly of the mountain hare (*L. timidus*) with the Alaskan hare (*L. othus*) (suggested by *ASTRAL* and *SVDquartets*; Supplementary Figs. S2a and S3a) and of the broom hare (*L. castroviejoi*) with the Corsican hare (*L. corsicanus*) (recovered with *ASTRAL*; Supplementary Fig. S2a). Both these cases involve sister taxa that likely share a very recent common ancestor and for which the classification as separate species has been debated (Alves et al. 2008; Melo-Ferreira et al. 2012). Also, we consistently estimated paraphyly of the Cape hare (*L. capensis*) with the Ethiopian hare (*L. fagani*), with the North African Cape hare specimen sharing a more recent common ancestor with Ethiopian hares than the South African Cape hare specimen (Supplementary Figs. S1, S2a and S3a). This may result from the known deep intraspecific divergences and paraphyly of Cape hare lineages (Lado et al. 2019), confirmed by our diversity and divergence estimates (Supplementary Tables S3 and S4).

Dating the species tree using either molecular-based dates extrapolated from deep fossil calibrations (Yamada et al. 2002; Matthee et al. 2004) or *Lepus* fossil record calibrations (Hibbard 1963; White 1991) recovered overlapping 95% high posterior density (HPD) intervals of divergence times for relatively recent splits (e.g., diversification of Eurasian and African species; Supplementary Table S5). However, *Lepus*-based fossil calibrations suggested more recent ages for deeper nodes (Supplementary Table S5 and Fig. S5). For instance, dates extrapolated from deeper fossil calibrations outside of *Lepus* sets the base of the hare radiation at ∼5.83 million year ago (Ma) (95% HPD 6.17-5.34 Ma), whereas *Lepus* fossil calibration recovered ∼4.05 Ma (95% HPD 5.00-3.18 Ma). All analyses were consistent in showing that the deepest branching events involved North American species: white-sided jackrabbit (*L. callotis*), black-tailed jackrabbit, and snowshoe hare (*L. americanus*) (Fig. 1b). In accordance, ancestral reconstruction of biogeographic distributions (Fig. 1c and Supplementary Table S6) supported an initial radiation of the genus in North America, with subsequent colonization of Eurasia (2.85 and 1.99 Ma for deeper and *Lepus*-fossil calibration, respectively) and Africa (1.85 and 1.33 Ma for deeper and fossil calibration, respectively). The North America distribution of white-tailed jackrabbits and Alaskan hares (Fig. 1a) likely results from one or two re-colonization events from Eurasia (Fig. 1c and Supplementary Table S6). These results also confirm the paraphyly of the jackrabbit species (Halanych et al. 1999).

### Incomplete Lineage Sorting and Introgression

We recovered a highly supported species trees across phylogenetic methods, albeit with extensive phylogenetic discordance among sequenced regions [average Robinson-Foulds (RF) pairwise distance between local trees was 0.73]. No local tree completely recovered the species tree topology (minimum RF distance between gene and species tree was 0.13) (Supplementary Fig. S6) and the majority rule consensus tree showed low internode certainty (Supplementary Fig. S7). Extensive phylogenetic discordance was also apparent in the *ASTRAL* species tree with all but three branches showing quartet scores below 0.6 (Fig. 1, Supplementary Figs. S2, S4 and Tables S7 and S8). Splits network analysis of individual gene trees also supported many alternative relationships (represented by cuboid structures connecting alternative topologies) particularly involving deeper branches (Fig. 2). We estimated similar levels of discordance between X-linked and autosomal genealogies and the inferred species trees (Supplementary Fig. S6). Furthermore, a species tree inferred with X-linked data differed from the genome-wide species tree and showed lower overall branch support (Supplementary Fig. S8 and Table S9). Measures of intraspecific diversity were relatively low within species [π between 0.13% (Corsican hare) and 0.63% (black-tailed jackrabbit] but overlapped in range with estimates of absolute genetic divergence between species [d_xy_ between 0.17% (broom hare and Corsican hare) and 1.11% (snowshoe hare and white-sided jackrabbit)] (Supplementary Tables S3 and S4). Some instances of low interspecific divergence reflect sister taxa where the species-level designations have been debated (e.g., the Corsican-Broom hare and Mountain-Alaskan hare complexes; Alves et al. 2008; Melo-Ferreira et al. 2012). Alternatively, high intraspecific diversity may reflect cryptic divergence within recognized species (e.g., snowshoe hares, Melo-Ferreira et al. 2014; Cape hares, Lado et al. 2019).

**Fig. 2.**
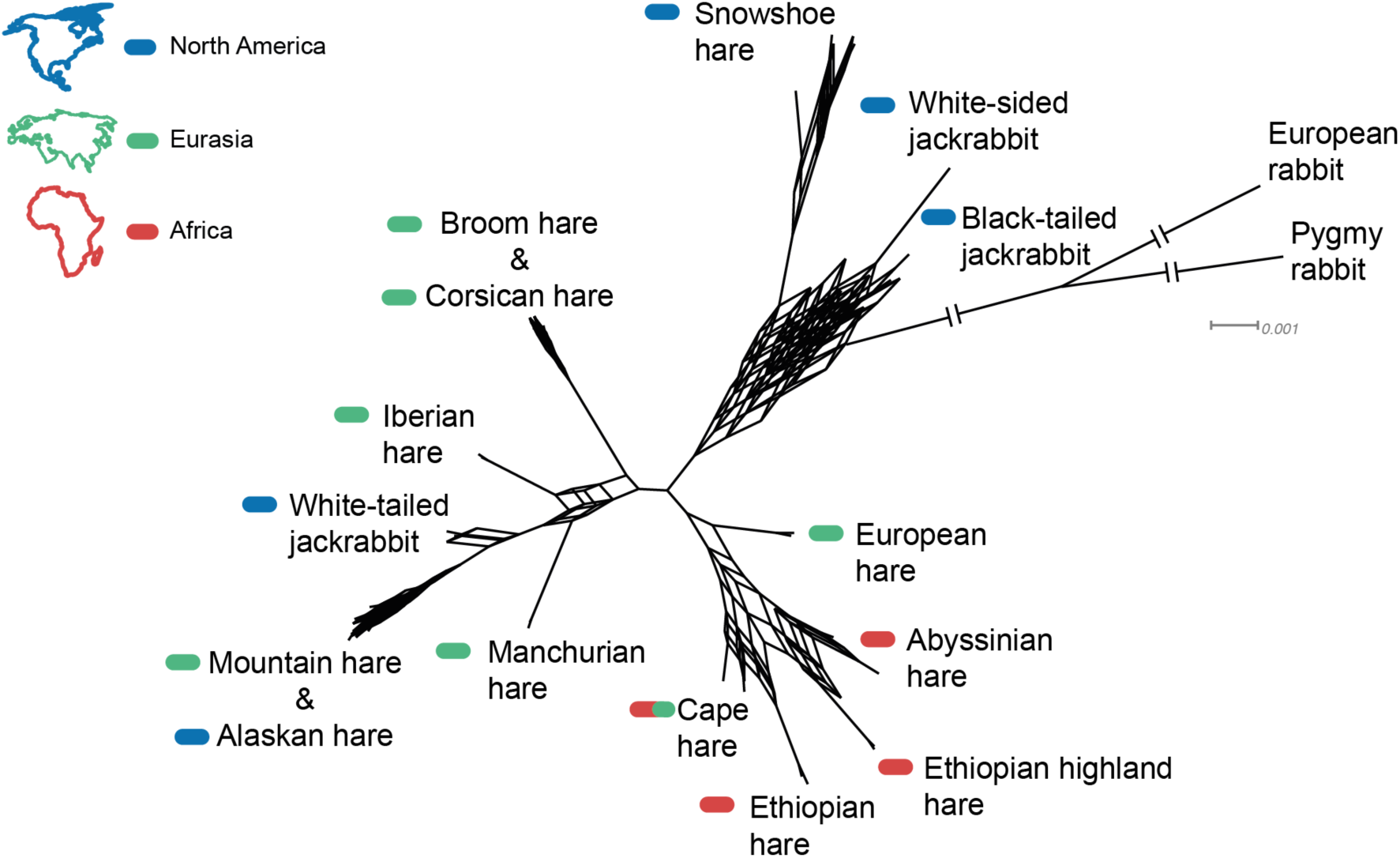
The hare (*Lepus* spp.) species tree is underlined by extensive gene tree incongruence. Split network constructed from 8889 gene trees (5% threshold) shows discordance among the gene tree topologies (cuboid structures represent alternative topologies) in deeper nodes of the species tree. Species are marked with colors corresponding to the continents where they are distributed.

On average, 49% of heterozygous sites in individuals of one species were shared with individuals from at least one other species (Supplementary Table S10). Such high levels of shared polymorphism between species coupled with extensive phylogenetic discordance across the genome could be explained by recent rapid speciation leading to extensive incomplete lineage sorting, secondary introgression between lineages, or a mixture of both processes. To differentiate these alternatives, we first tested if the proportions of shared-derived variants between species across the genome supported a strictly bifurcating history (expected with incomplete lineage sorting) by calculating minimum absolute D-statistics (D_min_) (Malinsky et al. 2018) for all trios of species in our dataset. We found that 88% of these comparisons were significantly different from zero (Bonferroni-corrected P < 0.05; Supplementary Fig. S9). This suggests that the level of shared alleles among the majority of species trios is incompatible with a single tree (even accounting for incomplete lineage sorting), providing overwhelming support for gene flow among species. This striking pattern of introgression could be driven by rampant gene flow among extant species pairs, or represent signatures of introgression between ancestral lineages that remain in modern species (Malinsky et al. 2018).

To disentangle temporally layered signatures of introgression, we combined analyses based on the multispecies network coalescent implemented in *PhyloNet* (Yu and Nakhleh 2015) (Supplementary Fig. S10), ancestral population graph reconstruction with *TreeMix* (based on 30,709 biallelic SNVs; Supplementary Figs. S11 and S12; Pickrell and Pritchard 2012), and estimates of admixture proportions among species based on the ‘f-branch’ metric (*f*_*b*_*(C)*) (Malinsky et al. 2018) (Fig. 3). Collectively, our results were consistent with recurrent gene flow between species layered across the diversification of *Lepus*. We detected introgression among extant species pairs within all of the major geographic regions that are currently sympatric, suggesting ongoing or recent hybridization (Fig. 3). For example, gene flow between black-tailed jackrabbits and snowshoe hares in North America (*f*_*b*_*(C) =* 19%; *P* = 2.04E-169), or between European brown hares and mountain hares from Eurasia (*f*_*b*_*(C)* = 18%; *P* = 1.73E-233; Fig. 3 and Supplementary Table S11). In general, we found decreased admixture proportions with increased genetic divergence between species, although this correlation was only significant when considering species with non-overlapping distributions (Fig. 4). Several species pairs with extant contact zones showed admixture even when genetic divergence was relatively high, such as snowshoe hares and black- tailed jackrabbits (d_xy_ = 0.97%; TMRCA ∼ 4.8 Ma, Supplementary Fig. S5 and Tables S4 and S5).

**Fig. 3.**
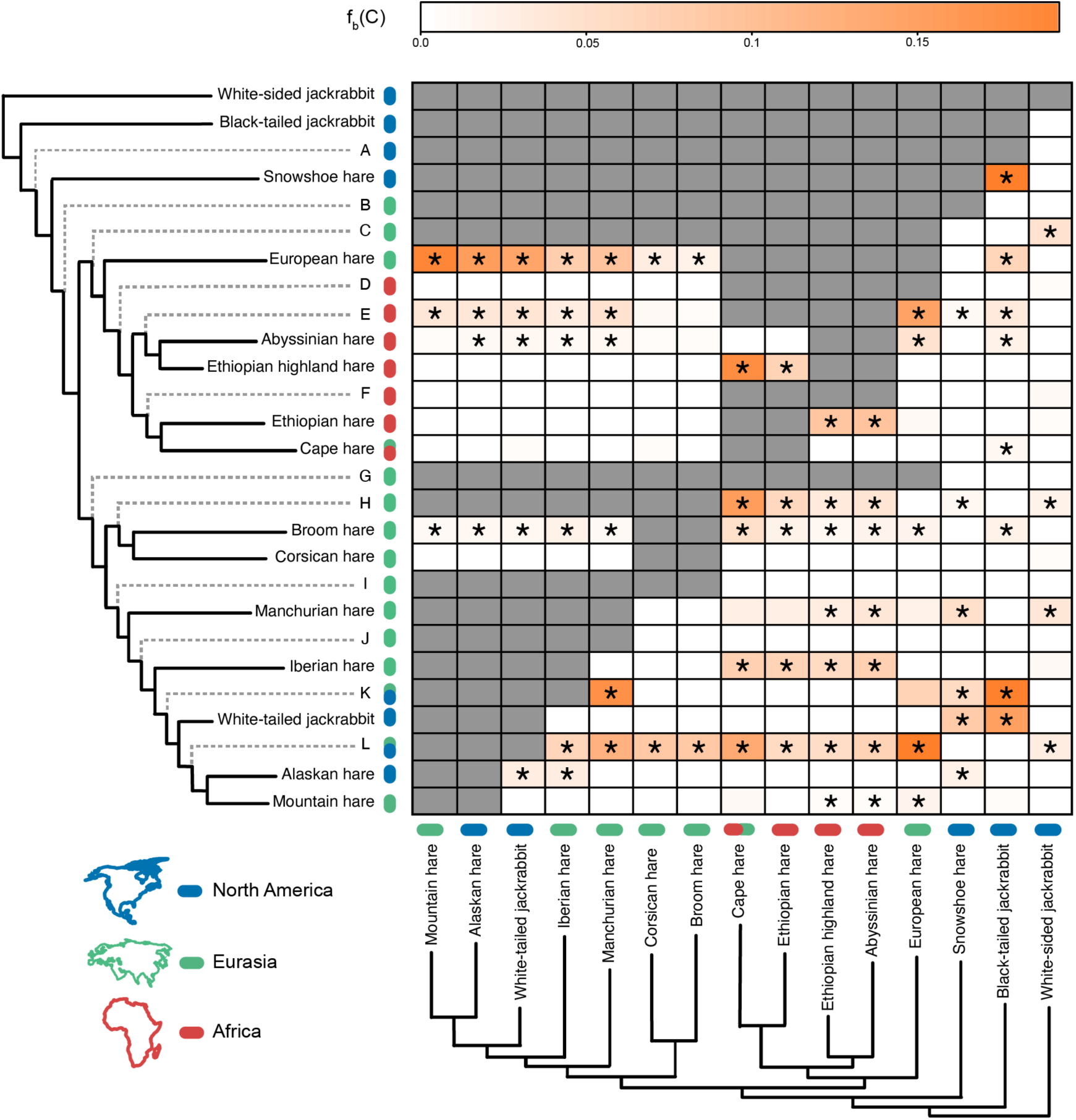
Admixture events are distributed across the hare (*Lepus* spp.) species tree. The branch-specific statistic *f*_*b*_*(C)* represents excess allele sharing between branches *b* (y-axis) and *C* (x-axis) of the species tree in Figure 1. The orange gradient represents the *f*_*b*_*(C)* score, gray represents tests not consistent with the species tree and asterisks denote block jackknifing significance at *P* ≤ 0.05 (after Bonferroni correction). Tips of the tree are colored according to their current distribution (two colors represent distribution in both areas), and ancestral tips (dashed gray and labeled with letters) are colored according to the ancestral reconstruction of distribution areas from Figure 1 (two colors represent equivocal inference of the two-character states). Ancestral tips are labeled from A to L corresponding to labels in Supplementary Table S11.

**Fig. 4.**
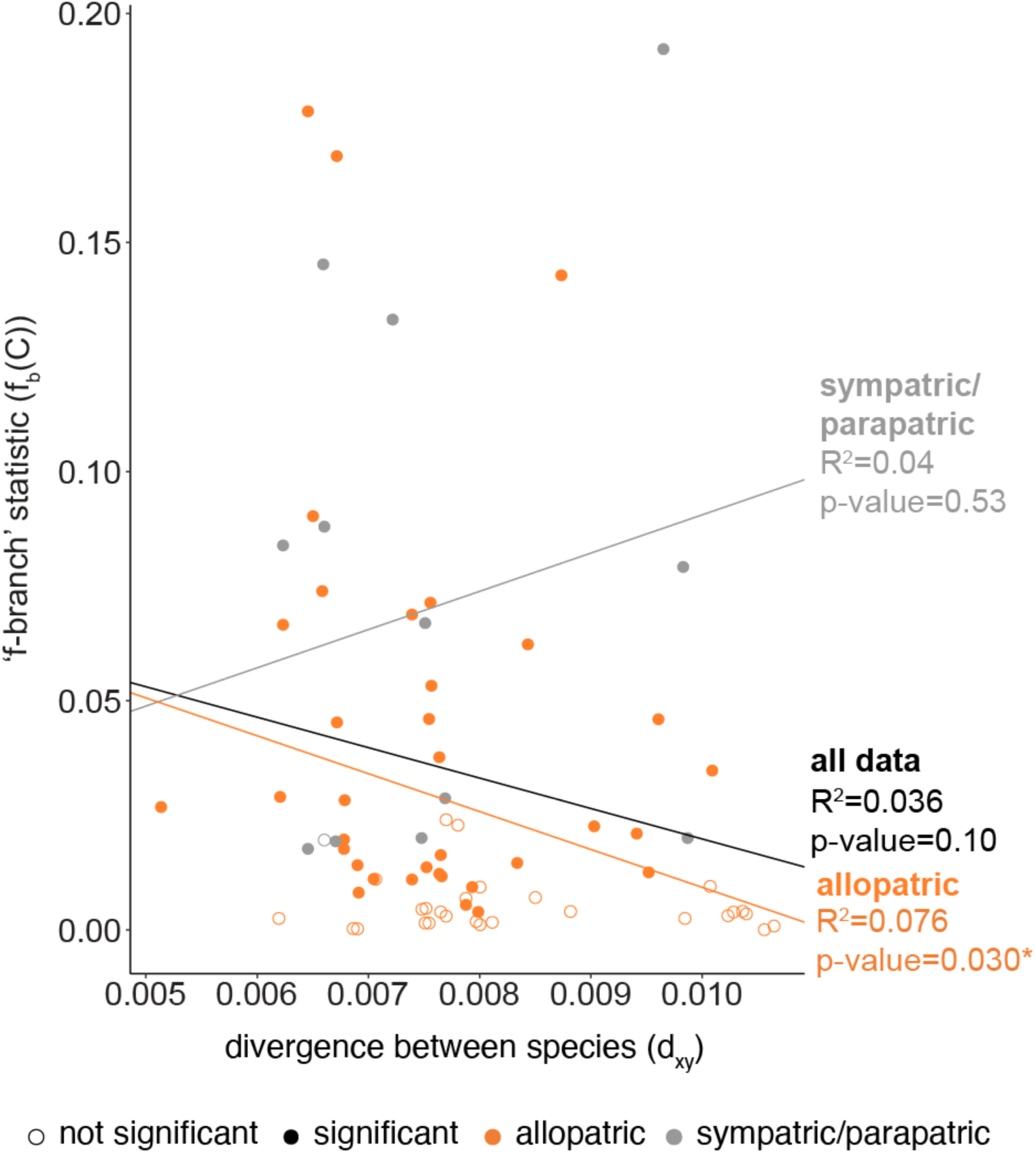
Admixture proportions decrease with genetic divergence between allopatric species. We plot *f*_*b*_*(C)* values against exome-wide divergence (d_xy_) for extant species pairs, coloring points when species have overlapping (sympatric/parapatric) or non-overlapping (allopatric) distributions. The tendency line represents a linear regression relating d_xy_ and *f*_*b*_*(C)* values calculated with function *lm()* in R.

We also found evidence for introgression between ancestral populations, which has affected deeper branches of the species tree. Network and *f*_*b*_*(C)* results suggest ancestral events of introgression connecting major diverged clades within Eurasia (e.g., European hares and the ancestor of the mountain hare/Alaskan hare/white-tailed jackrabbit clade), African and Eurasian lineages (e.g., ancestral of all African species and the Corsican and broom hare ancestral), and Eurasian and North American lineages (e.g., snowshoe hares and the ancestral lineage of the mountain hare/Alaskan hare/white-tailed jackrabbit; Fig. 3, 4, Supplementary Figs. S10 and S11, and Table S11). Finally, we found significant allele sharing among species that currently inhabit different continents, such as North American hares and species from Africa and Western Europe (Fig. 3 and 4). Trans-continental introgression was also suggested by a network with two reticulations and by ancestral population graph reconstruction (Supplementary Figs. S10c and S11). Given the inferred split times (Supplementary Fig. S5) and biogeography (Fig. 1) of this group, these results suggest that introgression affected the very early branches of the *Lepus* radiation, and that the genetic legacy of these gene flow events persists in the gene pool of descendant species today (Fig. 3 and 4).

### Genes affected by Ancestral Introgression

We detected ancestral introgression involving white-tailed jackrabbits, mountain hares, Alaskan hares, and snowshoe hares, which together have a circumpolar distribution across northern latitude habitats (Fig. 3 and Supplementary Fig. S10). These species inhabit the strongly seasonal boreal and/or temperate environments of North America and Eurasia (Smith et al. 2018), and have all evolved seasonal coat color molts from summer brown to winter white to maintain cryptic coloration with the onset of seasonal snow cover (Mills et al. 2018). Motivated by recent work showing that introgressive hybridization has shaped local adaptation in seasonal coat color changing species (Jones et al. 2018; Giska et al. 2019; Jones et al. 2020a), we next examined the contribution of gene flow between the ancestor of Alaskan hare/mountain hare/white-tailed jackrabbit and snowshoe hares (at least 0.98 Ma from lagomorph fossil calibrations and 0.71 Ma from *Lepus* fossil calibration; Supplementary Fig. S5 and Table S5) to standing variation in these species.

We performed fraction of admixture (*f*_*d*_) scans to detect genomic regions shared through ancestral introgression among these species. We detected 119 putative windows of ancient introgression across all major chromosomes, highlighting the genome-wide impact of ancestral introgression (Fig. 5; Supplementary Table S12). The *f*_*d*_ outlier windows of ancestral introgression contained 54 annotated genes (Supplementary Table S13). This set of genes was enriched for the gene ontology term “E-Box binding” (3 of the 54 genes; Supplementary Table S14), a DNA motif found in the promoters of many genes, suggesting that genomic regions affected by ancient introgression may be enriched for transcription factors involved in *trans*-regulation of gene expression. Among these transcription factors we found the circadian clock related gene *ARNTL2* (Sasaki et al. 2009) and pigmentation related gene *TCF4* (Furumura et al. 2001; Le Pape et al. 2009) (Fig. 5). In addition, the list of 54 genes includes a gene involved in brown fat differentiation (*EBF2*; Rajakumari et al. 2013), and a photoreceptor-related gene (*PDE6H*; Kohl et al. 2012).

**Fig. 5.**
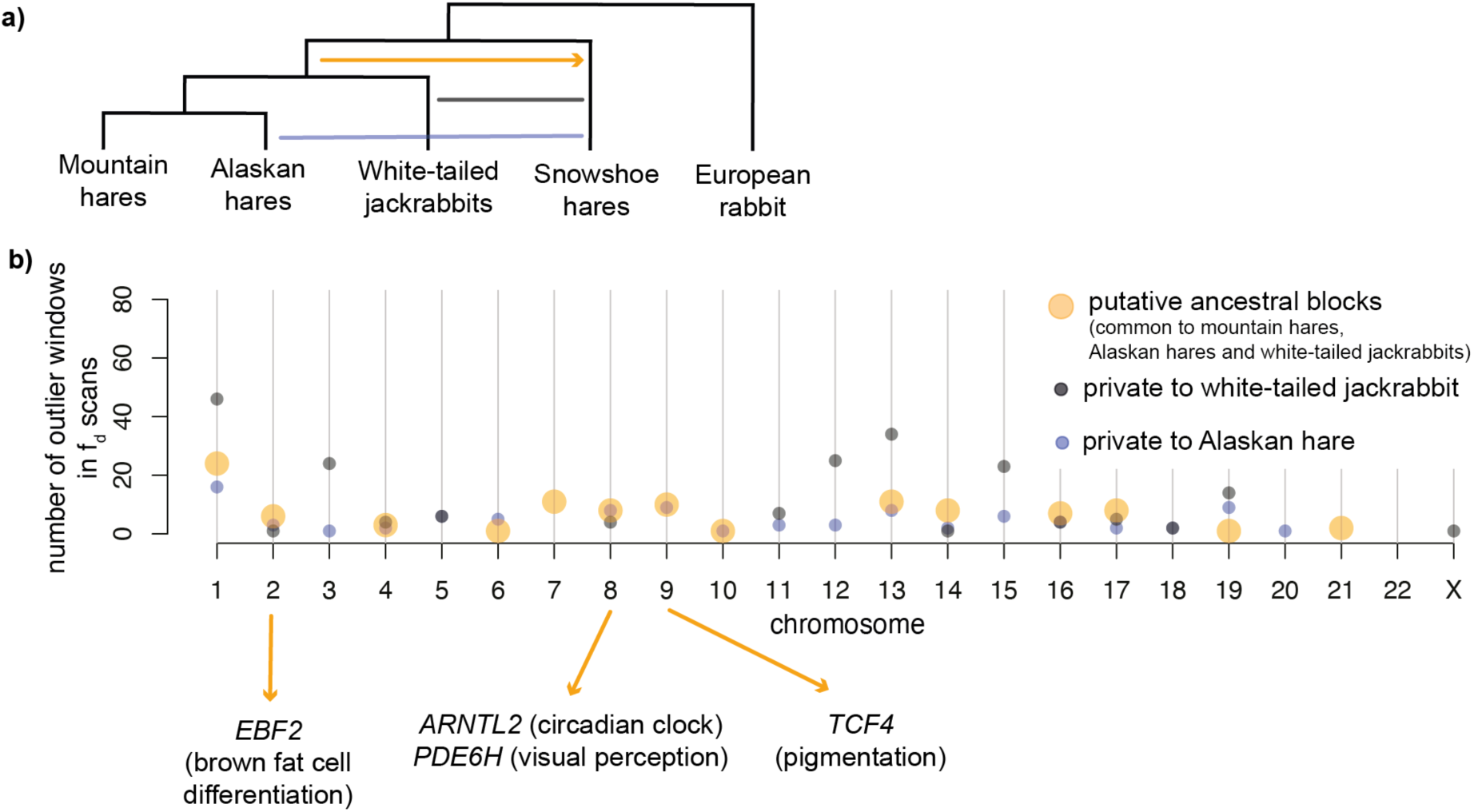
The impact of ancestral introgression on extant northern latitude species. **A)** Events of past and recent admixture inferred in this study involving snowshoe, Alaskan hares, mountain hares and white-tailed jackrabbits. The arrow indicates the direction of ancestral introgression inferred by *PhyloNet*; **B)** Genomic distribution of ancestral blocks of introgression inferred as shared outliers windows among fraction of admixture (*f*_*d*_) analysis testing for admixture among snowshoe hares and each one of the three other northern latitude species (orange), and blocks private to the analysis with Alaskan hares (blue) and white-tailed jackrabbits (black). Tests were run in 50 kb genomic sliding windows, and outlier windows are in the top 0.5% of the *f*_*d*_ distribution.

## Discussion

We used whole exome data and diverse phylogenetic analyses to tease apart signatures of stochastic lineage sorting and admixture across the evolutionary history of the *Lepus* radiation. By accounting for these sources of phylogenetic discordance, we were able to detect pervasive introgression between lineages that was layered across the evolution of this recent and rapid mammalian radiation. Below we discuss the biogeographic and evolutionary implications of our analyses, focusing on the long-term impacts of temporally layered hybridization in shaping patterns of shared genetic variation within and among extant species.

### The Effect of Persistent Gene Flow on Phylogenetic Inference

We present a resolved genome-wide phylogeny for the genus *Lepus* despite extensive incomplete lineage sorting and pervasive gene flow. Although we sampled just a subset of described *Lepus* species (15 of 32 species), our genome-wide experiment covered all major lineages across the worldwide range of hares resulting in the most extensive phylogeny of the genus to date, substantially extending previous analyses based on more limited genetic sampling (Halanych et al. 1999; Melo-Ferreira et al. 2012; Ge et al. 2013; Tolesa et al. 2017). A systematic evaluation of species limits and hare taxonomy is beyond the scope of our work as it would require genome-wide sequencing of an expanded inter and intraspecific sample. Our sampling design is however not expected to bias phylogenetic relationships of the sampled species and inference of discordance among deeper lineages, and may even underestimate species diversity, thus making our inferences of shared variation among species conservative.

To estimate species relationships, we combined species tree inferences that do not account for gene flow with network-based inferences that explicitly consider introgression. The resulting topologies were generally consistent between methods when considering up to three reticulations (Fig. 1 and Supplementary Fig. S10). However, there were clear limits to this approach. For example, the alternative placement of European, Corsican, and broom hares, as closer to the Eurasian or African *Lepus* clades depended on the number of reticulations considered (Supplementary Fig. S10). This uncertainty likely reflects long-term admixture among Eurasian and African lineages, including the European hare (Fig. 3 and Supplementary Fig. S10) whose range overlaps with species from both continents (Fig. 1a). Another example is the alternative sister relationship of snowshoe hares and black-tailed jackrabbits in the network with no reticulation (Supplementary Fig. S10), which could result from ancestral introgression between these species (see below). A network with four reticulations was less concordant than results assuming zero to three reticulations. While an increase number of reticulation events could better represent the widespread nature of gene flow uncovered in our work, allowing for more than three reticulations resulted in increased branch compression suggesting that the added instances of gene flow do not contribute to resolve gene-to-gene incongruences (Supplementary Fig. S10) (Yu and Nakhleh 2015; Wen et al. 2016).

Interestingly, we found that gene trees constructed from X-linked loci or autosome-linked loci showed similar levels of discordance with the whole-genome species tree (Supplementary Fig. S6). In general, the X chromosome could be expected to show less phylogenetic discordance due to a typically smaller effective population size (i.e., lower levels of incomplete lineage sorting assuming equal sex ratios) and a tendency to accumulate hybrid incompatibilities (i.e., the large X-effect) (Fontaine et al. 2015; Edelman et al. 2019; Li et al. 2019). Although there is evidence for reduced X-linked gene flow between some hybridizing European lineages (Seixas et al. 2018) very little is known about the genetic architecture of reproductive isolation between hare species. Moreover, higher gene tree/species tree concordance of loci involved in reproductive isolation may not be always expected, particularly when speciation events are clustered in time (Wang and Hahn 2018) as is the case in *Lepus* (Supplementary Fig. S5).

### The Timing and Biogeography of the Lepus Radiation

Overall, our results suggest that the *Lepus* diversification followed major climatic shifts that occurred during the late Miocene, Pliocene and Pleistocene, similar to other terrestrial mammals (Simpson 1947; Ge et al. 2013). We inferred that the genus originated in North America between 5.83 Ma (deeper fossil calibration) and 4.05 Ma (*Lepus* fossil calibration) (in general agreement with Hibbard 1963; Halanych et al. 1999; Melo-Ferreira et al. 2012; Ge et al. 2013). This inferred origin coincides with a global cold and dry period of the late Miocene that favored the expansion of grasslands worldwide (Osborne and Beerling 2006; Ge et al. 2013). The colonization of Eurasia around 2.77 Ma (deeper fossil calibration) or 1.99 Ma (*Lepus* fossil calibration) is also coincident with faunal exchanges between North America and Eurasia during late Pliocene and Pleistocene glacial periods that connected both continents through the Bering land bridge (Simpson 1947; Hopkins 1959; Cook et al. 2016). After colonizing Eurasia, *Lepus* entered Africa somewhere between the divergence of the European hare and the African ancestral lineage (1.78 Ma or 2.48 Ma from *Lepus* or deeper fossil calibrations respectively) and the divergence of all African species (1.33 Ma or 1.85 Ma from *Lepus* or deeper fossil calibrations respectively) (Supplementary Fig. S5 and Table S5).

Our biogeographic inferences confirm a secondary colonization of North America back from Eurasia likely within the last 1 Ma (Fig. 1 and Supplementary Fig. S5) (Halanych et al. 1999). The Eurasia-North America exchange of fauna during colder glacial periods of the Pleistocene is thought to have involved taxa adapted to colder environments (Simpson 1947; Hoberg et al. 2012). Consistent with this, we find that the *Lepus* re-colonization of North America involved the ancestral lineage of species now adapted to northern ecosystems (i.e., mountain hares, Alaskan hares, and white-tailed jackrabbits).

We note that our finding of pervasive introgression (Fig. 3, Supplementary Figs. S9, S10 and S11 and Table S11) may distort aspects of these biogeographic inferences (Leaché et al. 2014; Solís-Lemus et al. 2016; Long and Kubatko 2018; Li et al. 2019; Jiao et al. 2020). Assuming that the inferred structure of the *Lepus* phylogeny is correct, as discussed above, the general biogeographic reconstructions are probably robust. However, extensive ancestral gene flow is still likely to bias our estimates of species divergence (Leaché et al. 2014; Li et al. 2019). The lack of varied, independent, and reliable calibration points along our phylogeny hinders further evaluation of the impact of introgression on our time estimates, which therefore should be interpreted with caution.

### The Legacy of Introgression during the Rapid Radiation of Hares

Extensive reticulation across the *Lepus* phylogeny suggests that lineages often hybridized when they came into contact during their worldwide expansion. Some of the inferred reticulation events overlap with known recent introgression, such as between mountain hares and European hares (Levänen et al. 2018) or between snowshoe hares and black-tailed jackrabbits (Jones et al. 2018, 2020a). However, our analysis also reveals prevailing signatures of deeper hybridization between ancestral populations, suggesting a persistent contribution of secondary introgression during the diversification of the genus. These past hybridization events have resulted in extensive shared polymorphism among extant species (Fig. 3 and Supplementary Tables S10, S11 and Figs. S9, S10 and S11), with significant admixture still detected among species with non-overlapping distributions or inhabiting different continents (Fig. 3 and Fig. 4).

Our results highlight how ancient gene flow can obscure accurate detection of contemporary hybridization. Similar to other systems (Malinsky et al. 2018; Li et al. 2019; Edelman et al. 2019), we estimated exceedingly high levels of introgression based on D-statistics (e.g., 88% of D_min_ values across all possible species trios were significant), but much less reticulation when we took the phylogeny into account (e.g., 33% of f-branch statistics were significant). These findings suggest that some signatures of introgression between pairs of species may actually result from earlier hybridization between ancestral lineages. Thus, phylogenetic correlation causes non-independence of summary-statistics and can thus lead to false positive inferences of gene flow (Eaton et al. 2015; Malinsky et al. 2018; Li et al. 2019). Given these results, signatures of hybridization among closely related species should be interpreted in the context of broader phylogenetic relationships whenever possible.

We also detected some discrepancies between the magnitude of gene flow inferred here and in previous works, which underscores the challenges of quantifying introgression across a reticulating radiation. For instance, we inferred substantial overall admixture proportions between black-tailed jackrabbits and snowshoe hares (*f*_*b*_*(C)* = 19%, Fig. 3 and Supplementary Table S11). These estimates are one order of magnitude higher than recent studies suggesting that ∼2-3% of genomic variation in the Pacific Northwest snowshoe hare populations descends from a pulse of gene flow from black-tailed jackrabbits in the last ∼10,000 generations (Jones et al. 2018, 2020a). Ancient introgression persisting in all snowshoe hare populations could reconcile this discrepancy. Indeed, we recovered admixture in the same order of magnitude (*f*_hom_ ∼3%) when we infer admixture proportions among different snowshoe hare populations and black-tailed jackrabbits (Supplementary Table S15), which would then reflect recent and geographically localized introgression. However, our quantitative estimate of overall admixture proportions between these species (19%) also depends on the accurate reconstruction of the species tree, which in this instance involves a short internal branch that is not fully supported across methods, and a species (snowshoe hare) that has been involved in multiple instances of introgression with different hare lineages (Fig. 3, 5 and Supplementary Table S11 and Fig. S10).

Introgression between species is often limited by purifying selection against divergent alleles that are deleterious in hybrids (Schumer et al. 2018; Edelman et al. 2019), which agrees with the predominantly negative consequences of hybridization (Mayr 1963). Nonetheless, hybridization is also expected to produce novel allelic combinations that increase phenotypic variation (Grant and Grant 2019; Marques et al. 2019). If coincident with ecological opportunity, introgressed variation could broadly facilitate adaptation (Grant and Grant 2019; Taylor and Larson 2019). Recent work has shown at least two instances where introgression between *Lepus* species has driven local adaptation (Jones et al. 2018; Giska et al. 2019), and standing variation from introgression may have contributed even more generally to adaptation during the radiation. We found here that allelic combinations from ancient hybridization can be carried through several speciation events (Fig. 3 and 4), persisting in the gene pool of consecutive species due to the rapid succession of speciation events. Much of the large reservoir of shared variation may reflect stochastic sorting of neutral variation, while some could be maintained by positive selection (Guerrero and Hahn 2017), facilitating colonization of the diverse habitats currently inhabited by hare species, from desert to arctic environments (Ge et al. 2013; Smith et al. 2018).

In this respect, introgression between the ancestor of mountain hares/Alaskan hares/white-tailed jackrabbits and snowshoe hares (or an ancestral lineage) is particularly intriguing. These four species have adapted to highly seasonal environments through striking forms of phenotypic plasticity (e.g., seasonal coat color change; Mills et al. 2013; Zimova et al. 2018) that have been at least partially shaped by adaptive introgression at the *Agouti* pigmentation gene from non-color changing species (*ASIP*; Jones et al. 2018; Giska et al. 2019). Here we estimated that a pulse of ancient introgression occurred at least 0.71 Ma (*Lepus* fossil calibration; Supplementary Fig. S5) and affected genomic regions containing genes associated with circadian rhythm regulation (*ARNTL2*; Sasaki et al. 2009), pigmentation (*TCF4*; Furumura et al. 2001; Le Pape et al. 2009), thermoregulation (*EBF2*; Rajakumari et al. 2013) and visual perception (*PDE6H*; Kohl et al. 2012). While *ASIP* would also be a likely candidate for adaptive introgression between these lineages (Jones et al. 2018), our exome sequencing coverage of this region was too sparse for detailed window-based analysis.

The functions of these introgressed genes overlap with common physiological adaptations of northern latitude animals to seasonal conditions, such as higher metabolic rates, regulation of body temperature and non-shivering thermogenesis (Hart et al. 1965; Feist and Rosenmann 1975; Rogowitz 1990; Pyörnilä et al. 2008; Sheriff et al. 2009), seasonal camouflage (Grange 1932; Hewson 1958; Hansen and Bear 1963; Mills et al. 2018; Zimova et al. 2018), and visual acuity in response to dim winter light in northern latitudes (Stokkan et al. 2013). The functional relevance of these candidates to local adaptation must await further testing. Substantial introgression along the rapid diversification of a group of organisms, as we describe here, may bolster genetic variation within species and have a greater role in local adaptation than previously anticipated (Grant and Grant 2019; Taylor and Larson 2019). However, we also cannot exclude the possibility that some of these shared variants have been maintained by long-term balancing selection rather than secondary introgression (Supplementary Fig. S13; Smith and Kronforst 2013; Liu et al. 2015; Guerrero and Hahn 2017). Regardless of origin, the *Lepus* radiation provides an intriguing system by which to test the long-term evolutionary importance of shared genetic variation across a rapid radiation.

## Data Availability

Raw sequence reads were deposited at the Sequence Read Archive (SRA) under accession numbers SRR12020579 to SRR12020510 and BioProject PRJNA639005. Other data will be made available from the Dryad Digital Repository with link http://dx.doi.org/10.5061/dryad.[NNNN]. The pipeline and custom scripts used for this study are available at https://github.com/evochange.

## ACKNOWLEDGEMENTS

We thank Janet Rachlow, Gloria Portales, Armando Geraldes, José Carlos Brito, Jérôme Letty, Conrad Matthee, Alexei Kryukov, Ettore Randi, Christian Gortazar, Rafael Villafuerte, Fernando Ballesteros, Klaus Hackländer and Zbyszek Boratynski for assistance with sample collection. The Alaskan hare samples used in this work were generously provided by the University of Alaska Museum (voucher codes UAM42143 and UAM45545). We thank Fernando Seixas, Tiago Antão, Hannes Svardal, Nathaniel Edelman, the EVOCHANGE lab, the Good lab, and the UNVEIL network for helpful discussions on the generation and analysis of data. Funding and support for this research was provided by Fundação para a Ciência e a Tecnologia (FCT) (project grant “CHANGE” – PTDC/BIA-EVF/1624/2014, Portuguese National Funds) and the National Science Foundation (NSF) (EPSCoR OIA-1736249 and DEB-1907022 grant). MSF was supported by POPH-QREN funds from ESF and Portuguese MCTES/FCT (PD/BD/108131/2015 PhD grant in the scope of BIODIV PhD programme at Faculty of Sciences, University of Porto), Portuguese National Funds through FCT (PTDC/BIA-EVF/1624/2014), and by NSF (OIA-1736249). MRJ was supported by an NSF Graduate Research Fellowship (DGE-1313190). JM-F was supported by an FCT CEEC contract (CEECIND/00372/2018). Instrumentation, laboratory, and computational support was provided by CIBIO NEWGEN sequencing platform, supported by European Union’s Seventh Framework Program for research, technological development and demonstration under grant agreement no. 286431, by the University of Montana Genomics Core, supported by a grant from the M.J. Murdock Charitable Trust. A grant from the Eunice Kennedy Shriver National Institute of Child Health and Human Development (R01HD073439) supported the development of exome capture protocols that were utilized in the current study. Additional support was obtained from the Laboratoire International Associé (LIA) “Biodiversity and Evolution” funded by InEE (CNRS, France) and FCT (Portugal), COMPETE2020, PORTUGAL2020, and ERDF (POCI-01-0145-FEDER-022184), and from Portugal- United States of America Research Networks Program funds from Fundação Luso-Americana para o Desenvolvimento (FLAD) to PCA and MSF.

## REFERENCES

Alves P.C., Melo-Ferreira J., Branco M., Suchentrunk F., Ferrand N., Harris D.J. 2008. Evidence for genetic similarity of two allopatric European hares (*Lepus corsicanus* and *L. castroviejoi*) inferred from nuclear DNA sequences. Mol. Phylogenet. Evol. 46:1191–7.

Árnason Ú., Lammers F., Kumar V., Nilsson M.A., Janke A. 2018. Whole-genome sequencing of the blue whale and other rorquals finds signatures for introgressive gene flow. Sci. Adv. 4.

Van der Auwera G.A., Carneiro M.O., Hartl C., Poplin R., del Angel G., Levy-Moonshine A., Jordan T., Shakir K., Roazen D., Thibault J., Banks E., Garimella K. V, Altshuler D., Gabriel S., DePristo M.A. 2013. From fastQ data to high-confidence variant calls: The genome analysis toolkit best practices pipeline. Curr. Protoc. Bioinforma. 11.

Barlow A., Cahill J.A., Hartmann S., Theunert C., Xenikoudakis G., Fortes G.G., Paijmans J.L.A., Rabeder G., Frischauf C., Grandal-d’Anglade A., García-Vázquez A., Murtskhvaladze M., Saarma U., Anijalg P., Skrbinšek T., Bertorelle G., Gasparian B., Bar-Oz G., Pinhasi R., Slatkin M., Dalén L., Shapiro B., Hofreiter M. 2018. Partial genomic survival of cave bears in living brown bears. Nat. Ecol. Evol. 2:1563–1570.

Barth J.M.I., Gubili C., Matschiner M., Tørresen O.K., Watanabe S., Egger B., Han Y.S., Feunteun E., Sommaruga R., Jehle R., Schabetsberger R. 2020. Stable species boundaries despite ten million years of hybridization in tropical eels. Nat. Commun. 11:1–13.

Bennett D.J., Sutton M.D., Turvey S.T. 2017. Treeman: An R package for efficient and intuitive manipulation of phylogenetic trees. BMC Res. Notes. 10:30.

Borowiec M.L. 2016. AMAS: A fast tool for alignment manipulation and computing of summary statistics. PeerJ. 2016:e1660.

Carneiro M., Rubin C.J., Palma F. Di, Albert F.W., Alföldi J., Barrio A.M., Pielberg G., Rafati N., Sayyab S., Turner-Maier J., Younis S., Afonso S., Aken B., Alves J.M., Barrell D., Bolet G., Boucher S., Burbano H.A., Campos R., Chang J.L., Duranthon V., Fontanesi L., Garreau H., Heiman D., Johnson J., Mage R.G., Peng Z., Queney G., Rogel-Gaillard C., Ruffier M., Searle S., Villafuerte R., Xiong A., Young S., Forsberg-Nilsson K., Good J.M., Lander E.S., Ferrand N., Lindblad-Toh K., Andersson L. 2014. Rabbit genome analysis reveals a polygenic basis for phenotypic change during domestication. Science. 345:1074–1079.

Chifman J., Kubatko L. 2014. Quartet inference from SNP data under the coalescent model. Bioinformatics. 30:3317–3324.

Cook J.A., Galbreath K.E., Bell K.C., Campbell M.L., Carrière S., Colella J.P., Dawson N.G., Dunnum J.L., Eckerlin R.P., Fedorov V., Greiman S.E., Haas G.M., Haukisalmi V., Henttonen H., Hope A.G., Jackson D., Jung T.S., Koehler A. V, Kinsella J.M., Krejsa D., Kutz S.J., Liphardt S., MacDonald S.O., Malaney J.L., Makarikov A., Martin J., McLean B.S., Mulders R., Nyamsuren B., Talbot S.L., Tkach V. V, Tsvetkova A., Toman H.M., Waltari E.C., Whitman J.S., Hoberg E.P., Cook J., Bell K., Colella J., Jackson D., Krejsa D., Liphardt S., McLean B., Galbreath K., Haas G., Toman H., Campbell M., Dunnum J., MacDonald S., Carrière S., Mulders R., Dawson N., Fedorov V., Greiman S., Jung T., Koehler A., Kinsella J., Kutz S., Malaney J., Makarikov A., Martin J., Nyamsuren B., Waltari Aaron E., Hoberg Animal E. 2016. The Beringian Coevolution Project: holistic collections of mammals and associated parasites reveal novel perspectives on evolutionary and environmental change in the North. Arct. Sci. 3:585–617.

Degnan J.H. 2018. Modeling Hybridization Under the Network Multispecies Coalescent. Syst. Biol. 67:786–799.

Durinck S., Moreau Y., Kasprzyk A., Davis S., De Moor B., Brazma A., Huber W. 2005. BioMart and Bioconductor: a powerful link between biological databases and microarray data analysis. Bioinformatics. 21:3439–3440.

Durinck S., Spellman P.T., Birney E., Huber W. 2009. Mapping identifiers for the integration of genomic datasets with the R/ Bioconductor package biomaRt. Nat. Protoc. 4:1184–1191.

Eaton D.A.R., Hipp A.L., González-Rodríguez A., Cavender-Bares J. 2015. Historical introgression among the American live oaks and the comparative nature of tests for introgression. Evolution. 69:2587–2601.

Edelman N.B., Frandsen P.B., Miyagi M., Clavijo B., Davey J., Dikow R.B., García-Accinelli G., Van Belleghem S.M., Patterson N., Neafsey D.E., Challis R., Kumar S., Moreira G.R.P., Salazar C., Chouteau M., Counterman B.A., Papa R., Blaxter M., Reed R.D., Dasmahapatra K.K., Kronforst M., Joron M., Jiggins C.D., Owen McMillan W., Palma F. Di, Blumberg A.J., Wakeley J., Jaffe D., Mallet J. 2019. Genomic architecture and introgression shape a butterfly radiation. Science. 366:594–599.

Feist D.D., Rosenmann M. 1975. Seasonal sympatho-adrenal and metabolic responses to cold in the Alaskan snowshoe hare (*Lepus americanus macfarlani*). Comp. Biochem. Physiol. Part A Physiol. 51:449–455.

Fontaine M.C., Pease J.B., Steele A., Waterhouse R.M., Neafsey D.E., Sharakhov I. V, Jiang X., Hall A.B., Catteruccia F., Kakani E., Mitchell S.N., Wu Y.-C., Smith H.A., Love R.R., Lawniczak M.K., Slotman M.A., Emrich S.J., Hahn M.W., Besansky N.J. 2015. Extensive introgression in a malaria vector species complex revealed by phylogenomics. Science. 347:1258522–1258522.

Furumura M., Potterf S.B., Toyofuku K., Matsunaga J., Muller J., Hearing V.J. 2001. Involvement of ITF2 in the transcriptional regulation of melanogenic genes. J. Biol. Chem. 276:28147–54.

Ge D., Wen Z., Xia L., Zhang Z., Erbajeva M., Huang C., Yang Q. 2013. Evolutionary History of Lagomorphs in Response to Global Environmental Change. PLoS One. 8:e59668.

Giska I., Farelo L., Pimenta J., Seixas F.A., Ferreira M.S., Marques J.P., Miranda I., Letty J., Jenny H., Hackländer K., Magnussen E., Melo-Ferreira J. 2019. Introgression drives repeated evolution of winter coat color polymorphism in hares. Proc. Natl. Acad. Sci. 116:24150–24156.

Gittelman R.M., Schraiber J.G., Vernot B., Mikacenic C., Wurfel M.M., Akey J.M. 2016. Archaic Hominin Admixture Facilitated Adaptation to Out-of-Africa Environments. Curr. Biol. 26:1–8.

Grange W.B. 1932. The Pelages and Color Changes of the Snowshoe Hare, *Lepus americanus phaeonotus*, Allen. J. Mammal. 13:99.

Grant P.R., Grant B.R. 2019. Hybridization increases population variation during adaptive radiation. Proc. Natl. Acad. Sci. 116:23216–23224.

Green R.E., Krause J., Briggs A.W., Maricic T., Stenzel U., Kircher M., Patterson N., Li H., Zhai W., Fritz M.H.-Y., Hansen N.F., Durand E.Y., Malaspinas A.-S., Jensen J.D., Marques-Bonet T., Alkan C., Prüfer K., Meyer M., Burbano H.A., Good J.M., Schultz R., Aximu-Petri A., Butthof A., Höber B., Höffner B., Siegemund M., Weihmann A., Nusbaum C., Lander E.S., Russ C., Novod N., Affourtit J., Egholm M., Verna C., Rudan P., Brajkovic D., Kucan Ž., Gušic I., Doronichev V.B., Golovanova L. V, Lalueza-Fox C., de la Rasilla M., Fortea J., Rosas A., Schmitz R.W., Johnson P.L.F., Eichler E.E., Falush D., Birney E., Mullikin J.C., Slatkin M., Nielsen R., Kelso J., Lachmann M., Reich D., Pääbo S. 2010. A draft sequence of the Neandertal genome. Science. 328:710–722.

Guerrero R.F., Hahn M.W. 2017. Speciation as a sieve for ancestral polymorphism. Mol. Ecol. 26:5362–5368.

Halanych K.M., Demboski J.R., van Vuuren B.J., Klein D.R., Cook J. a. 1999. Cytochrome b phylogeny of North American hares and jackrabbits (*Lepus*, Lagomorpha) and the effects of saturation in outgroup taxa. Mol. Phylogenet. Evol. 11:213–221.

Hansen R.M., Bear G.D. 1963. Winter coats of white-tailed jackrabbits in southwestern Colorado. J. Mammology. 44:420–422.

Hart J.S., Pohl H., Tener J.S. 1965. Seasonal acclimatization in varying hare (*Lepus americanus*). Can. J. Zool. 43:731–744.

Hewson R. 1958. Moults and winter whitening in the mountain hare *Lepus timidus scoticus*, Hilzheimer. Proc. Zool. Soc. London. 131:99–108.

Hibbard C. 1963. The Origin of the P 3 Pattern of *Sylvilagus, Caprolagus, Oryctolagus* and *Lepus*. J. Mammal. 44:1–15.

Hoberg E.P., Galbreath K.E., Cook J.A., Kutz S.J., Polley L. 2012. Northern Host-Parasite Assemblages. History and Biogeography on the Borderlands of Episodic Climate and Environmental Transition. Advances in Parasitology. Academic Press. p. 1–97.

Hopkins D.M. 1959. Cenozoic history of the Bering land bridge. Science. 129:1519–1528.

Huson D.H., Bryant D. 2006. Application of phylogenetic networks in evolutionary studies. Mol. Biol. Evol. 23:254–267.

Inoue J., Donoghue P.C.J., Yang Z. 2010. The Impact of the Representation of Fossil Calibrations on Bayesian Estimation of Species Divergence Times. Syst. Biol. 59:74–89.

Jiao X., Flouri T., Rannala B., Yang A.Z. 2020. The Impact of Cross-Species Gene Flow on Species Tree Estimation. Syst. Biol. 0:1–18.

Jones M.R., Mills L.S., Alves P.C., Callahan C.M., Alves J.M., Lafferty D.J.R., Jiggins F.M., Jensen J.D., Melo-Ferreira J., Good J.M. 2018. Adaptive introgression underlies polymorphic seasonal camouflage in snowshoe hares. Science. 360:1355–1358.

Jones M.R., Mills L.S., Jensen J.D., Good J.M. 2020a. The origin and spread of locally adaptive seasonal camouflage in snowshoe hares. Am. Nat. https://doi.org/10.1086/710022

Jones M.R., Mills L.S., Jensen J.D., Good J.M. 2020b. Convergent evolution of seasonal camouflage in response to reduced snow cover across the snowshoe hare range. Evolution. Early View. https://doi.org/10.1111/evo.13976

Kinoshita G., Nunome M., Kryukov A.P., Kartavtseva I. V., Han S.-H., Yamada F., Suzuki H. 2019. Contrasting phylogeographic histories between the continent and islands of East Asia: Massive mitochondrial introgression and long-term isolation of hares (Lagomorpha: *Lepus*). Mol. Phylogenet. Evol. 136:65–75.

Kohl S., Coppieters F., Meire F., Schaich S., Roosing S., Brennenstuhl C., Bolz S., van Genderen M.M., Riemslag F.C.C., Lukowski R., den Hollander A.I., Cremers F.P.M., De Baere E., Hoyng C.B., Wissinger B. 2012. A Nonsense Mutation in *PDE6H* Causes Autosomal-Recessive Incomplete Achromatopsia. Am. J. Hum. Genet. 91:527–532.

Kozak K.M., McMillan O., Joron M., Jiggins C.D. 2018. Genome-wide admixture is common across the *Heliconius* radiation. bioRxiv. https://doi.org/10.1101/414201

Lado S., Alves P.C., Islam M.Z., Brito J.C., Melo-Ferreira J. 2019. The evolutionary history of the Cape hare (*Lepus capensis sensu lato*): insights for systematics and biogeography. Heredity. 123:634–646.

Lamichhaney S., Berglund J., Almén M.S., Maqbool K., Grabherr M., Martinez-Barrio A., Promerová M., Rubin C.-J., Wang C., Zamani N., Grant B.R., Grant P.R., Webster M.T., Andersson L. 2015. Evolution of Darwin’s finches and their beaks revealed by genome sequencing. Nature. 518:371–375.

Leaché A.D., Harris R.B., Rannala B., Yang Z. 2014. The Influence of Gene Flow on Species Tree Estimation: A Simulation Study. Syst. Biol. 63:17–30.

Levänen R., Thulin C.-G., Spong G., Pohjoismäki J.L.O. 2018. Widespread introgression of mountain hare genes into Fennoscandian brown hare populations. PLoS One. 13:e0191790.

Lewis P.O. 2001. A Likelihood Approach to Estimating Phylogeny from Discrete Morphological Character Data. Syst. Biol. 50:913–925.

Li G., Figueiró H. V, Eizirik E., Murphy W.J., Yoder A. 2019. Recombination-Aware Phylogenomics Reveals the Structured Genomic Landscape of Hybridizing Cat Species. Mol. Biol. Evol. 36:2111–2126.

Li H. 2011. A statistical framework for SNP calling, mutation discovery, association mapping and population genetical parameter estimation from sequencing data. Bioinformatics. 27:2987–2993.

Li H. 2013. Aligning sequence reads, clone sequences and assembly contigs with BWA-MEM. arXiv Prepr. 1303.

Li H., Handsaker B., Wysoker A., Fennell T., Ruan J., Homer N., Marth G., Abecasis G., Durbin R. 2009. The Sequence Alignment/Map format and SAMtools. Bioinformatics. 25:2078–9.

Liu J., Yu L., Arnold M.L., Wu C.-H., Wu S.-F., Lu X., Zhang Y.-P. 2011. Reticulate evolution: frequent introgressive hybridization among chinese hares (genus *Lepus*) revealed by analyses of multiple mitochondrial and nuclear DNA loci. BMC Evol. Biol. 11:223.

Liu K.J., Steinberg E., Yozzo A., Song Y., Kohn M.H., Nakhleh L. 2015. Interspecific introgressive origin of genomic diversity in the house mouse. 112:196–201.

Long C., Kubatko L. 2018. The Effect of Gene Flow on Coalescent-based Species-Tree Inference. Syst. Biol. 67:770–785.

Lopez-Martinez N. 2008. The lagomorph fossil record and the origin of the European rabbit. In: Alves P.C., Ferrand N., Häcklander K., editors. Lagomorph Biology: Evolution, Ecology and Conservation. Verlag Berlin Heidelberg: Springer. p. 27–46.

Maddison W.P., Maddison D.R. 2018. Mesquite: a modular system for evolutionary analysis. Version 3.51.

Malinsky M., Svardal H., Tyers A.M., Miska E.A., Genner M.J., Turner G.F., Durbin R. 2018. Whole-genome sequences of Malawi cichlids reveal multiple radiations interconnected by gene flow. Nat. Ecol. Evol. 2:1940–1955.

Marques D.A., Meier J.I., Seehausen O. 2019. A Combinatorial View on Speciation and Adaptive Radiation. Trends Ecol. Evol. 34:531–544.

Martin S.H., Davey J.W., Jiggins C.D. 2015. Evaluating the use of ABBA-BABA statistics to locate introgressed loci. Mol. Biol. Evol. 32:244–257.

Matthee C., Van Vuuren B., Bell D., Robinson T. 2004. A Molecular Supermatrix of the Rabbits and Hares (Leporidae) Allows for the Identification of Five Intercontinental Exchanges During the Miocene. Syst. Biol. 53:433–447.

Mayr E. 1963. Animal species and evolution. Eugen. Rev. 55:226–228.

Meier J.I., Marques D.A., Mwaiko S., Wagner C.E., Excoffier L., Seehausen O. 2017. Ancient hybridization fuels rapid cichlid fish adaptive radiations. Nat. Commun. 8:14363.

Melo-Ferreira J., Alves P.C. 2018. Systematics of lagomorphs. In: Smith A., Alves P.C., Häcklander K., editors. Lagomorphs: pikas, rabbits, and hares of the World. John Hopkins University Press. p. 9–12.

Melo-Ferreira J., Boursot P., Carneiro M., Esteves P.J., Farelo L., Alves P.C. 2012. Recurrent introgression of mitochondrial DNA among hares (*Lepus* spp.) revealed by species-tree inference and coalescent simulations. Syst. Biol. 61:367–381.

Melo-Ferreira J., Seixas F.A., Cheng E., Mills L.S., Alves P.C. 2014. The hidden history of the snowshoe hare, *Lepus americanus*: extensive mitochondrial DNA introgression inferred from multilocus genetic variation. Mol. Ecol.:4617–4630.

Meyer M., Kircher M. 2010. Illumina Sequencing Library Preparation for Highly Multiplexed Target Capture and Sequencing. Cold Spring Harb. Protoc. 2010.

Mills L.S., Bragina E., Kumar A. V., Zimova M., Feltner J., Davis B., Lafferty D., Häcklander K., Alves P.C., Good J.M., Melo-Ferreira J., Abramov A., Lopatina N., Fay K. 2018. Winter coat color polymorphisms identify global hotspots for evolutionary rescue from climate change. Science. 359:1033–1036.

Mills L.S., Zimova M., Oyler J., Running S., Abatzoglou J.T., Lukacs P.M. 2013. Camouflage mismatch in seasonal coat color due to decreased snow duration. Proc. Natl. Acad. Sci. U. S. A. 110:7360–7365.

Osborne C.P., Beerling D.J. 2006. Nature’s green revolution: the remarkable evolutionary rise of C 4 plants. Philos. Trans. R. Soc. B Biol. Sci. 361:173–194.

Pagel M. 1999. The maximum likelihood approach to reconstructing ancestral character states of discrete characters on phylogenies. Syst. Biol. 48:612–622.

Le Pape E., Passeron T., Giubellino A., Valencia J.C., Wolber R., Hearing V.J. 2009. Microarray analysis sheds light on the dedifferentiating role of agouti signal protein in murine melanocytes via the Mc1r. Proc. Natl. Acad. Sci. U. S. A. 106:1802–1807.

Paradis E., Claude J., Strimmer K. 2004. APE: Analyses of Phylogenetics and Evolution in R language. Bioinformatics. 20:289–290.

Pease J.B., Hahn M.W. 2015. Detection and Polarization of Introgression in a Five-Taxon Phylogeny. Syst. Biol. 64:651–662.

Pickrell J.K., Pritchard J.K. 2012. Inference of Population Splits and Mixtures from Genome-Wide Allele Frequency Data. PLoS Genet. 8:e1002967.

Pyörnilä A., Putaala A., Hissa R., Sulkava S. 2008. Adaptations to environment in the mountain hare (*Lepus timidus*): thermal physiology and histochemical properties of locomotory muscles. Can. J. Zool. 70:1325–1330.

Rajakumari S., Wu J., Ishibashi J., Lim H.-W., Giang A.-H., Won K.-J., Reed R.R., Seale P. 2013. EBF2 Determines and Maintains Brown Adipocyte Identity. Cell Metab. 17:562–574.

Raudvere U., Kolberg L., Kuzmin I., Arak T., Adler P., Peterson H., Vilo J. 2019. g:Profiler: a web server for functional enrichment analysis and conversions of gene lists (2019 update). Nucleic Acids Res. 47:191–198.

Reich D., Thangaraj K., Patterson N., Price A.L., Singh L. 2009. Reconstructing Indian population history. Nature. 461:489–494.

dos Reis M., Yang Z. 2011. Approximate Likelihood Calculation on a Phylogeny for Bayesian Estimation of Divergence Times. Mol. Biol. Evol. 28:2161–2172.

dos Reis M., Yang Z. 2019. Bayesian Molecular Clock Dating Using Genome-Scale Datasets. In: Anisimova M., editor. Evolutionary Genomics. New York, NY: Springer New York. p. 309–330.

Rogowitz G.L. 1990. Seasonal Energetics of the White-Tailed Jackrabbit (*Lepus townsendii*). J. Mammal. 71:277–285.

Salichos L., Stamatakis A., Rokas A. 2014. Novel information theory-based measures for quantifying incongruence among phylogenetic trees. Mol. Biol. Evol. 31:1261–1271.

Sambrook E., Fritsch F., Maniatis T. 1989. Molecular cloning. Cold Spring Harbour, NY: Cold Spring Harbour Press.

Sarver B.A.J., Keeble S., Cosart T., Tucker P.K., Dean M.D., Good J.M. 2017. Phylogenomic Insights into Mouse Evolution Using a Pseudoreference Approach. Genome Biol. Evol. 9:726–739.

Sasaki M., Yoshitane H., Du N.-H., Okano T., Fukada Y. 2009. Preferential Inhibition of BMAL2-CLOCK Activity by PER2 Reemphasizes Its Negative Role and a Positive Role of BMAL2 in the Circadian Transcription. J. Biol. Chem. 284:25149–25159.

Sayyari E., Whitfield J.B., Mirarab S. 2018. DiscoVista: Interpretable visualizations of gene tree discordance. Mol. Phylogenet. Evol. 122:110–115.

Schliep K.P. 2011. phangorn: phylogenetic analysis in R. Bioinform. 27:592–593.

Schluter D., Price T., Mooers A.O., Ludwig D. 1997. Likelihood of ancestral states in adaptive radiation. Evolution. 51:1699–1711.

Schumer M., Xu C., Powell D.L., Durvasula A., Skov L., Holland C., Blazier J.C., Sankararaman S., Andolfatto P., Rosenthal G.G., Przeworski M. 2018. Natural selection interacts with recombination to shape the evolution of hybrid genomes. Science. 360:656–660.

Seixas F.A., Boursot P., Melo-Ferreira J. 2018. The genomic impact of historical hybridization with massive mitochondrial DNA introgression. Genome Biol. 19:91.

Sheriff M.J., Kuchel L., Boutin S., Humphries M.M. 2009. Seasonal Metabolic Acclimatization in a Northern Population of Free-Ranging Snowshoe Hares, *Lepus americanus*. J. Mammal. 90:761–767.

Simpson G.G. 1947. Holarctic mammalian faunas and continental relationships during the cenozoic. Bull. Geol. Soc. Am. 58:613–688.

Smith A.T., Johnston C.H., Alves P.C., Häcklander K. 2018. Lagomorphs: Pikas, Rabbits and Hares of the World. Baltimore: Johns Hopkins University Press.

Smith J., Kronforst M.R. 2013. Do *Heliconius* butterfly species exchange mimicry alleles? Biol. Lett. 9:20130503–20130503.

Solís-Lemus C., Bastide P., Ané C. 2017. PhyloNetworks: A Package for Phylogenetic Networks. Mol. Biol. Evol. 34:3292–3298.

Solís-Lemus C., Yang M., Ané C. 2016. Inconsistency of Species Tree Methods under Gene Flow. Syst. Biol. 65:843–851.

Stamatakis A. 2018. RAxML version 8: a tool for phylogenetic analysis and post–analysis of large phylogenies. Bioinformatics. 30:1312–1313.

Stokkan K.-A., Folkow L., Dukes J., Neveu M., Hogg C., Siefken S., Dakin S.C., Jeffery G. 2013. Shifting mirrors: adaptive changes in retinal reflections to winter darkness in Arctic reindeer. Proc. R. Soc. B Biol. Sci. 280:20132451.

Svardal H., Quah F.X., Malinsky M., Ngatunga B.P., Miska E.A., Salzburger W., Genner M.J., Turner G.F., Durbin R. 2020. Ancestral Hybridization Facilitated Species Diversification in the Lake Malawi Cichlid Fish Adaptive Radiation. Mol. Biol. Evol. 37:1100–1113.

Swofford D.L. 2003. PAUP*. Phylogenetic Analysis Using Parsimony (* and Other Methods).

Tange O. 2011. GNU Parallel - The Command-Line Power Tool.

Taylor S.A., Larson E.L. 2019. Insights from genomes into the evolutionary importance and prevalence of hybridization in nature. Nat. Ecol. Evol. 3:170–177.

Than C., Degnan J.H., Nakhleh L. 2011. Coalescent Histories on Phylogenetic Networks and Detection of Hybridization Despite Incomplete Lineage Sorting. Syst. Biol. 60:138–149.

Tolesa Z., Bekele E., Tesfaye K., Ben Slimen H., Valqui J., Getahun A., Hartl G.B., Suchentrunk F. 2017. Mitochondrial and nuclear DNA reveals reticulate evolution in hares (*Lepus* spp., Lagomorpha, Mammalia) from Ethiopia. PLoS One. 12:e0180137.

Vanderpool D., Minh B.Q., Lanfear R., Hughes D., Murali S., Harris R.A., Raveendran M., Muzny D.M., Gibbs R.A., Worley K.C., Rogers J., Hahn M.W. 2020. Primate phylogenomics uncovers multiple rapid radiatio. bioRxiv. https://doi.org/10.1101/2020.04.15.043786

Wang R.J., Hahn M.W. 2018. Speciation genes are more likely to have discordant gene trees. Evol. Lett. 2:281–296.

Wen D., Yu Y., Hahn M.W., Nakhleh L. 2016. Reticulate evolutionary history and extensive introgression in mosquito species revealed by phylogenetic network analysis. Mol. Ecol. 25:2361–2372.

White J.A. 1991. North American Leporinae (Mammalia: Lagomorpha) from late Miocene (Clarendonian) to latest Pliocene (Blancan). J. Vertebr. Paleontol. 11:67–89.

Yamada F., Takaki M., Suzuki H. 2002. Molecular phylogeny of Japanese Leporidae, the Amami rabbit *Pentalagus furnessi*, the Japanese hare *Lepus brachyurus*, and the mountain hare *Lepus timidus*, inferred from mitochondrial DNA sequences. Genes Genet. Syst. 77:107–16.

Yang Z. 2007. PAML 4: Phylogenetic analysis by maximum likelihood. Mol. Biol. Evol. 24:1586–1591.

Yu Y., Degnan J.H., Nakhleh L. 2012. The Probability of a Gene Tree Topology within a Phylogenetic Network with Applications to Hybridization Detection. PLoS Genet. 8:1002660.

Yu Y., Dong J., Liu K.J., Nakhleh L. 2014. Maximum likelihood inference of reticulate evolutionary histories. Proc. Natl. Acad. Sci. 111:16448–16453.

Yu Y., Nakhleh L. 2015. A maximum pseudo-likelihood approach for phylogenetic networks. BMC Genomics. 16:S10.

Zhang C., Rabiee M., Sayyari E., Mirarab S. 2018. ASTRAL-III: Polynomial time species tree reconstruction from partially resolved gene trees. BMC Bioinformatics. 19:153.

Zimova M., Hackländer K., Good J.M., Melo-Ferreira J., Alves P.C., Mills L.S. 2018. Function and underlying mechanisms of seasonal colour moulting in mammals and birds: what keeps them changing in a warming world? Biol. Rev. 93:1478–1498.

