## Appendix for "The legacy of recurrent introgression during the radiation of hares"

### EXTENDED METHODS

*Exome Capture Design and Laboratory Work*

Genomic DNA was extracted using either a saline extraction method (Sambrook et al. 1989) or the DNeasy Blood & Tissue Kit (Qiagen) (Supplementary Table S1) and quantified using a Qubit fluorometer (Invitrogren). For exome capture, we used the capture design and followed the exome library generation protocol described in Jones et al. (2018). The capture probe set was designed to target all protein-coding and untranslated regions (UTRs) annotated in the European rabbit reference genome (OryCun2.0) and novel UTRs inferred from previously published snowshoe hares transcriptomes (Ferreira et al. 2017). The capture probe set consisted of 213,164 probes covering 61.73 Megabases (Mb), of which ~25 Mb are protein-coding regions, ~28 Mb are untranslated regions and ~9 Mb are intron/intergenic regions. Library preparation steps followed a modified version of Meyer and Kircher (2010). Genomic DNA was sheared to ~300 base pairs (bps) with a Covaris E220evolution ultrasonicator, followed by cleaning using homemade 1.2X carboxyl-coated magnetic beads (Rohland and Reich 2012). Blunt-end repair, adapter ligation and adapter fill-in were done according to Meyer and Kircher (2010). To avoid over-amplification, we determined the number of cycles for dual-indexing PCR by performing quantitative PCR using a DyNamo Flash SYBR Green qPCR kit (Thermo Scientific) and a Stratagene Mx3000P thermocycler (Applied Biosystems). Dual-indexing PCR was performed in 50 µl Herculase II fusion polymerase and 3.5µl library template. The PCR thermal conditions consisted of an initial denaturation at 98ºC for 2 minutes, followed by 10-16 cycles of 98ºC for 20 seconds, and 60ºC for 20 seconds, and 72ºC for 20 seconds, with a final elongation step at 72ºC for 5 minutes. We quantified individually indexed libraries using an Implen P330 NanoPhotometer or Qubit (Invitrogen Qubit Quantitation system LTI) and visualized on an Agilent 2230 TapeStation. Then, we performed hybridization reactions following NimbleGen SeqCap EZ v.4.3 protocol in two separate sets of equimolarly pooled indexed libraries (Supplementary Table S1) together with samples not included in this work, for a total of 31 and 29 libraries in each pool. The 72-hour hybridization reaction included 5μg of $C_0t$-1 DNA isolated from snowshoe hare liver to prevent capture of repetitive regions. After hybridization, we washed and recovered hybridized DNA fragments following the NimbleGen protocol (Roche). We performed a post-capture PCR for each captured library pool with 1X Herculase II reaction buffer, 200 μM each dNTP, 2μM each primer, 1ul Herculase II fusion polymerase and 20μl library template. The thermal profile of the PCR consisted of a 45 second denaturation step at 95ºC, 16 cycles of 98ºC for 15 seconds, 60ºC for 30 seconds and 72ºC for 30 seconds, followed by a final elongation step at 72ºC for 5 minutes. The amplified libraries were cleaned with 1.8X AMPure XP beads (Agencourt). Enrichment of the capture reaction was evaluated with qPCR using a DyNAmo FLASH SYBR Green qPCR kit with both on-target and off-target primer sets. The two pools were each sequenced in two lanes of an Illumina HiSeq1500 sequencer (125 bp paired-end reads) at CIBIO-InBIO's New-Gen sequencing platform, Portugal.

*Genotype Calling and Filtering*

We mapped the data of all individuals (including those used to generate a pseudo-reference) back to each species-specific pseudo-reference using *bwa mem* (v.0.7.12-r1039; Li 2013) with default options. We created and sorted resulting bams using *samtools* (1.4; Li et al. 2009). We then added read groups (*AddOrReplaceReadGroups*), removed duplicates (*MarkDuplicates*) and performed indel realignment (*RealignerTargetCreator* and *IndelRealigner*) using *Picard* (v1.140; http://broadinstitute.github.io/picard/). We calculated coverage statistics and capture efficiency using *Picard*'s *CalculateHSMetrics* and called variants individually using the *bcftools* (1.4; Li 2011) mpileup, call and filter pipeline. We excluded indels and filtered genotypes with mapping quality smaller than 20 ($MQ < 20$), with phred based quality smaller than 20 (QUAL < 20), depth smaller than 6 (DP < 6), with depth more than 3 times higher than the average coverage for that individual (Supplementary Table S2), less than 10 bases from an indel (--SnpGap 10), and with genotype quality smaller than 20 if homozygous derived or heterozygous (GQ <= 20). With a custom script (available at https://github.com/evochange), we constructed consensus genome sequences for each individual, where we included all genotype positions passing filter and all other genomic positions were hard masked. We opted to call genome-wide sequences to facilitate the generation of input files for the various analysis. Furthermore, given all genomes were in the same coordinate system as the rabbit reference genome (OryCun2.0), we could quickly access annotations for any given region.

*Generating alignments for Species Tree Inference*

To generate a maximum likelihood tree for the concatenated whole exome, we retained positions with information for all individuals (no missing data allowed) from whole-genome alignments using *TriSeq* (1.0.1rc2), from *TriFusion* (http://odiogosilva.github.io/TriFusion/).

For the coalescent species tree analysis using *ASTRAL*, we generated 50 kilobase (kb) windows with no overlap from the individual genome consensus sequences using *msa_split* (--windows 50000,0) from *phast* (1.4) (http://compgen.cshl.edu/phast/). Then, to determine which windows contained regions included in the exome capture, we used *bedtools* (1.9; Quinlan and Hall 2010) *makewindows* to produce a bed file with the genomic windows' coordinates and we excluded windows not included in targeted exome regions extended by 200 bp. Furthermore, for each alignment, we filtered positions with missing data for more than 30% of the individuals using *TriSeq* and, finally, excluded alignments shorter than 1000 bps. Then, we estimated a tree for each window alignment with *RAxML* (v8.2.10; Stamatakis 2018) using a simultaneous maximum likelihood (ML) search and 100 rapid bootstraps, under the GTR+Γ model of sequence evolution (-f a option), and setting the European rabbit and pygmy rabbit as outgroups. Prior to the *ASTRAL* analysis, we unrooted the gene trees using R package *ape* (Paradis et al. 2004). For each resultant gene tree, we used the bootstrap trees generated by *RAxML* to calculate a majority rule consensus tree and calculate tree certainty scores using *RAxML* (-L MRE option) (Salichos et al. 2014).

We also estimated a coalescent species tree with single nucleotide variants (SNVs) with *SVDquartets* (Chifman and Kubatko 2014). Using genome-wide sequences, we produced an alignment containing only *Lepus* species, and used *snp-sites* (v2.3.3; Page et al. 2016) to extract a vcf file containing SNVs in the alignment. We filtered positions in this vcf with missing data for more than 30% of the individuals using a custom script (*filter_missing_vcf.py*; available at https://github.com/evochange). Furthermore, we filtered out positions not included in the targeted regions extended by 200 bp, using *bedtools intersect*. We recovered SNVs distanced 10 kb along the genome using a custom script (*filter_bed.py*; available at https://github.com/evochange) to respect the assumption of independence among sites. With the resulting filtered SNV bed file, we produced a SNV fasta alignment containing 45,779 SNVs for all *Lepus* individuals, European rabbit and pygmy rabbit using *bedtools getfasta*. This multi species alignment was converted to nexus with *AMAS* (option *convert*) (Borowiec 2016).

*Generating alignments for Bayesian Time Inference*

To extract the coding sequence (CDS) of all genes included in our capture (18,798 genes) we selected the longest transcript per gene using *biomaRt* (v2.34.2; Durinck et al. 2005, 2009) package in R. Then, we extracted the coding exons using *bedtools getfasta* along with a gff3 file for each gene from the rabbit ENSEMBL 94 database (OryCun2.0). We concatenated the coding exons using custom scripts (available at https://github.com/evochange). We filtered alignments with more than 20% missing data as calculated by *AMAS summary*, which resulted in 9015 CDS alignments. With these, we constructed a concatenated alignment in phylip format with three partitions corresponding to the three codon positions using *extract_123_CDS_algn.py* (available at https://github.com/evochange), *AMAS concat* and *AMAS convert.*

*Generating SNV alignments for TreeMix*

To perform ancestral population graph reconstruction with *TreeMix* (Pickrell and Pritchard 2012), our pipeline was similar to the *SVDquartets* analysis, except that we only included *Lepus* species and used the white-sided jackrabbit as outgroup. We used a custom script (*fake_phase_fasta_algn.py* available at https://github.com/evochange) to randomly phase the SNV alignment for each chromosome. Then, we used the script *TreeMix_from_nex.py* (https://github.com/mgharvey) to convert phased fasta alignments to the *TreeMix* input, assigning all alleles from a species to a single population to calculate allele counts. This script excluded all multi-allelic sites.

*Nucleotide diversity, Genetic Divergence and Admixture Proportion*

To calculate several variants of the D-statistics ($D_{min}$, $f_b(C)$ and $f_d$) in our dataset, genetic divergence ($d_{xy}$) and nucleotide diversity ($\pi$), we used or based our analysis in the *genomics general* collection of scripts and tutorials by Simon Martin (https://github.com/simonhmartin/; last accessed January 14, 2019). All scripts mentioned in this section are from *genomics general,* unless otherwise stated. We started by converting an alignment of the genomic consensus sequences for all *Lepus* individuals and the European rabbit reference to a geno file (we included all genomic positions to preserve genomic coordinates) using the *seq2Geno.py* script. Using *filterGenotypes.py*, we filtered genotypes with missing data for more than 30% of the individuals.

— *Genetic Divergence and Nucleotide Diversity*

With the script *popgenWindows.py*, we calculated nucleotide diversity ($\pi$) per species and pairwise genetic distance between species ($d_{xy}$) in non-overlapping sliding windows of 1 Mb with more than 100 sites and averaged across all windows to obtain an exome wide estimate per species.

— *The minimum D-statistics ($D_{min}$)*

For $D_{min}$, we calculated derived allele frequencies for all species using *freq.py*, grouping all individuals of a species using a population file (--popFile) and fixing the European rabbit reference as the outgroup. With a custom R script (*D-stat.R* available at https://github.com/evochange), we calculated D (ABBA-BABA; Green et al. 2010) for all combinations of possible trios of species in our dataset (1365 calculations). To calculate z-scores, we performed block jackknifing using *jackknife.R*, establishing blocks of 1 Mb and removing blocks with missing data. D values with Bonferroni-corrected $P \leq 0.05$ were considered significantly different from zero.

— *The 'f-branch' statistic*

The 'f-branch' statistic ($f_b(C)$) requires the calculation of 'admixture proportion' ($f_G$) among all species pairs (Martin et al. 2015; Malinsky et al. 2018). Therefore, in addition to calculating allele frequencies for all species as described for $D_{min}$, and to produce P3a and P3b frequencies, we 1) calculated allele frequencies for all individuals per species with two samples, 2) calculated allele frequencies splitting species sampling in two populations for species with more than two samples or; 3) split the geno file entry for the single white-sided jackrabbit in two haploid chromosomes with a custom script (*make_haploid_geno.py* available at https://github.com/evochange), and used the two resulting entries to calculate two sets of allele frequencies, following Martin et al. (2015). With a custom script (*fd_tree_calculations.R* available at https://github.com/evochange) that implements the R package *treeman* (1.1.3; (Bennett et al. 2017), we calculated the necessary taxa combinations for $f_b(C)$ based on the *ASTRAL* species tree (Fig. 1), following Malinsky et al. (2018). Then, we calculated $f_G$ for each combination of taxa (using *f_G.R* available at https://github.com/evochange), calculating z-scores for all combinations following what is described for $D_{min}$. The exception was the white-sided jackrabbit, for which we randomly shuffled the alleles in each position for every jackknife iteration to avoid biases in phasing. We finally calculated $f_b(C)$ for each branch *b* and species *C* as described by Malinsky *et al.*, (2018) (*fb_C.R* available at https://github.com/evochange)

— *The fraction of admixture ($f_d$)*

We performed three $f_d$ scans with the conformation ((Iberian hare, P2), Snowshoe hare), where P2 is either Alaskan hare, mountain hare or white-tailed jackrabbit. We used *ABBABABAwindows.py* to calculate $f_d$ in sliding windows of 50 kb (5 kb step and minimum of 100 sites) and defined outliers as the top 0.5% $f_d$ values in each test. We inspected the functions of genes contained in the outlier windows, by annotating them using *biomaRt* package in R and the OryCun2.0 reference, and performing an enrichment analysis in *g:Profiler* (accessed September 2019; Raudvere et al. 2019) with the default parameters. We also compared the average $d_{xy}$ for $f_d$ outlier windows against the exome-wide $d_{xy}$ distribution between each pair of P2 and P3 species calculation $d_{xy}$ in windows of 50 kb, with 5 kb step for windows with more than 2000 sites, using *popgenWindows.py*.

### References

Bennett D.J., Sutton M.D., Turvey S.T. 2017. Treeman: An R package for efficient and intuitive manipulation of phylogenetic trees. BMC Res. Notes. 10:30.

Borowiec M.L. 2016. AMAS: A fast tool for alignment manipulation and computing of summary statistics. PeerJ. 2016:e1660.

Chifman J., Kubatko L. 2014. Quartet inference from SNP data under the coalescent model. Bioinformatics. 30:3317–3324.

Durinck S., Moreau Y., Kasprzyk A., Davis S., De Moor B., Brazma A., Huber W. 2005. BioMart and Bioconductor: a powerful link between biological databases and microarray data analysis. Bioinformatics. 21:3439–3440.

Durinck S., Spellman P.T., Birney E., Huber W. 2009. Mapping identifiers for the integration of genomic datasets with the R/ Bioconductor package biomaRt. Nat. Protoc. 4:1184–1191.

Ferreira M.S., Alves P.C., Callahan C.M., Marques J.P., Mills L.S., Good J.M., Melo-Ferreira J. 2017. The transcriptional landscape of seasonal coat color molt in the snowshoe hare. Mol. Ecol. 26:4173–4185.

Green R.E., Krause J., Briggs A.W., Maricic T., Stenzel U., Kircher M., Patterson N., Li H., Zhai W., Fritz M.H.-Y., Hansen N.F., Durand E.Y., Malaspinas A.-S., Jensen J.D., Marques-Bonet T., Alkan C., Prüfer K., Meyer M., Burbano H.A., Good J.M., Schultz R., Aximu-Petri A., Butthof A., Höber B., Höffner B., Siegemund M., Weihmann A., Nusbaum C., Lander E.S., Russ C., Novod N., Affourtit J., Egholm M., Verna C., Rudan P., Brajkovic D., Kucan Ž., Gušic I., Doronichev V.B., Golovanova L. V, Lalueza-Fox C., de la Rasilla M., Fortea J., Rosas A., Schmitz R.W., Johnson P.L.F., Eichler E.E., Falush D., Birney E., Mullikin J.C., Slatkin M., Nielsen R., Kelso J., Lachmann M.,

Reich D., Pääbo S. 2010. A draft sequence of the Neandertal genome. Science. 328:710–722.

Jones M.R., Mills L.S., Alves P.C., Callahan C.M., Alves J.M., Lafferty D.J.R., Jiggins F.M., Jensen J.D., Melo-Ferreira J., Good J.M. 2018. Adaptive introgression underlies polymorphic seasonal camouflage in snowshoe hares. Science. 360:1355–1358.

Li H. 2011. A statistical framework for SNP calling, mutation discovery, association mapping and population genetical parameter estimation from sequencing data. Bioinformatics. 27:2987–2993.

Li H. 2013. Aligning sequence reads, clone sequences and assembly contigs with BWA-MEM. 00:1–3.

Li H., Handsaker B., Wysoker A., Fennell T., Ruan J., Homer N., Marth G., Abecasis G., Durbin R. 2009. The Sequence Alignment/Map format and SAMtools. Bioinformatics. 25:2078–9.

Malinsky M., Svardal H., Tyers A.M., Miska E.A., Genner M.J., Turner G.F., Durbin R. 2018. Whole-genome sequences of Malawi cichlids reveal multiple radiations interconnected by gene flow. Nat. Ecol. Evol. 2:1940–1955.

Martin S.H., Davey J.W., Jiggins C.D. 2015. Evaluating the use of ABBA-BABA statistics to locate introgressed loci. Mol. Biol. Evol. 32:244–257.

Meyer M., Kircher M. 2010. Illumina Sequencing Library Preparation for Highly Multiplexed Target Capture and Sequencing. Cold Spring Harb. Protoc. 2010.

Page A.J., Taylor B., Delaney A.J., Soares J., Seemann T., Keane J.A., Harris S.R. 2016. SNP-sites: rapid efficient extraction of SNPs from multi-FASTA alignments. Microb. genomics. 2:e000056.

Paradis E., Claude J., Strimmer K. 2004. APE: Analyses of Phylogenetics and Evolution in R language. Bioinformatics. 20:289–290.

Pickrell J.K., Pritchard J.K. 2012. Inference of Population Splits and Mixtures from Genome-Wide Allele Frequency Data. PLoS Genet. 8:e1002967.

Quinlan A.R., Hall I.M. 2010. BEDTools: a flexible suite of utilities for comparing genomic features. Bioinformatics. 26:841–842.

Raudvere U., Kolberg L., Kuzmin I., Arak T., Adler P., Peterson H., Vilo J. 2019. g:Profiler: a web server for functional enrichment analysis and conversions of gene lists (2019 update). Nucleic Acids Res. 47:191–198.

Rohland N., Reich D. 2012. Cost-effective, high-throughput DNA sequencing libraries for multiplexed target capture. Genome Res. 22:939–946.

Salichos L., Stamatakis A., Rokas A. 2014. Novel information theory-based measures for quantifying incongruence among phylogenetic trees. Mol. Biol. Evol. 31:1261–1271.

Sambrook E., Fritsch F., Maniatis T. 1989. Molecular cloning. Cold Spring Harbour, NY: Cold Spring Harbour Press.

Stamatakis A. 2018. RAxML version 8: a tool for phylogenetic analysis and post-analysis of large phylogenies. Bioinformatics. 30:1312–1313.
