## Supplementary Figures for "The legacy of recurrent introgression during the radiation of hares"

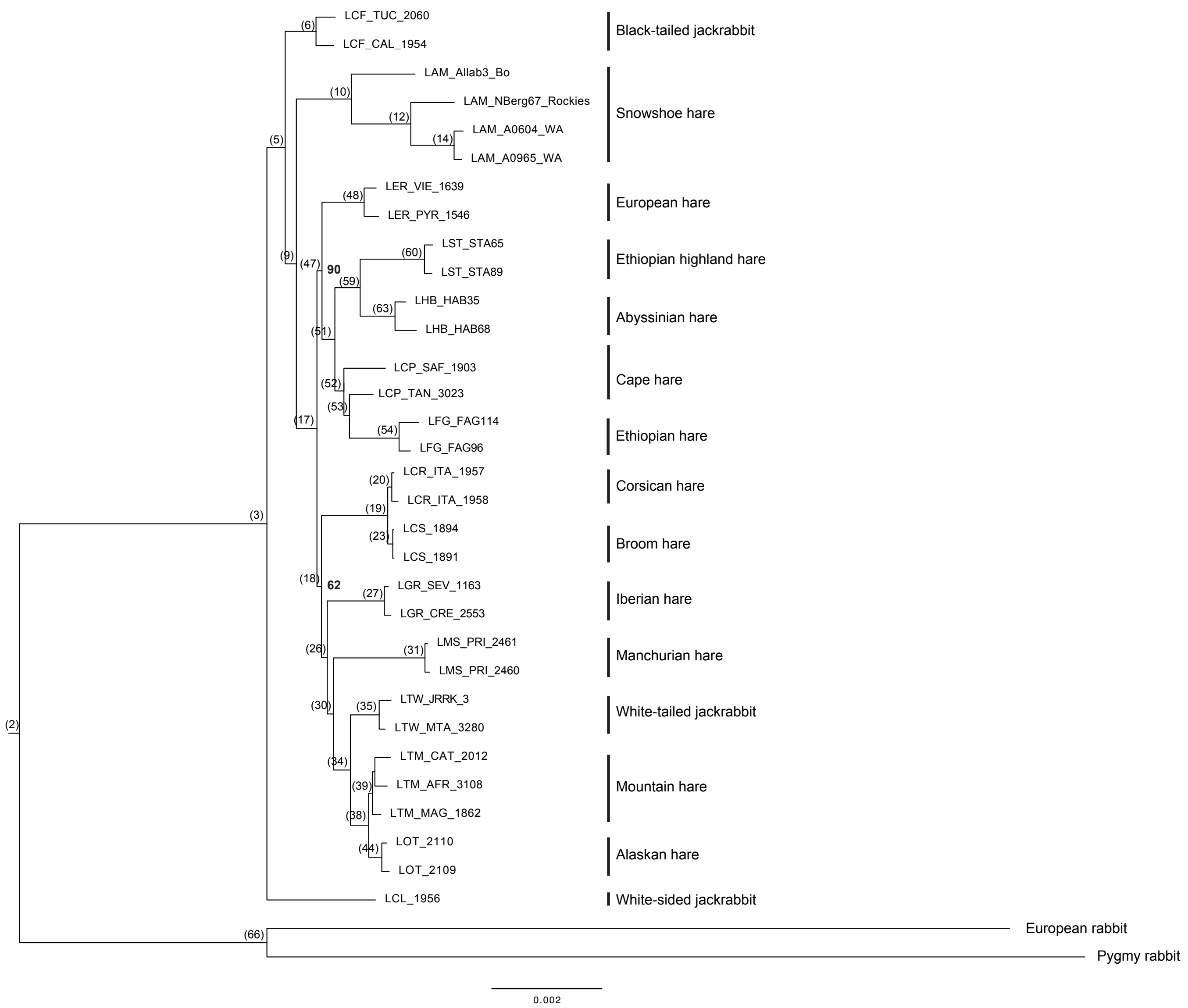


Figure S1 – Maximum likelihood tree of 11,949,529 base pair exome alignment (positions with no missing data for any individual), generated using a simultaneous maximum likelihood (ML) search in *RAxML* and rapid bootstrapping run under the GTR+Γ model of sequence evolution (autoMRE option), and setting the European and pygmy rabbits as outgroups. We show branch supports for branches with support smaller than 100 in bold. Node labels in parenthesis correspond to node labels in the ancestral reconstruction output in Supplementary Table S6.

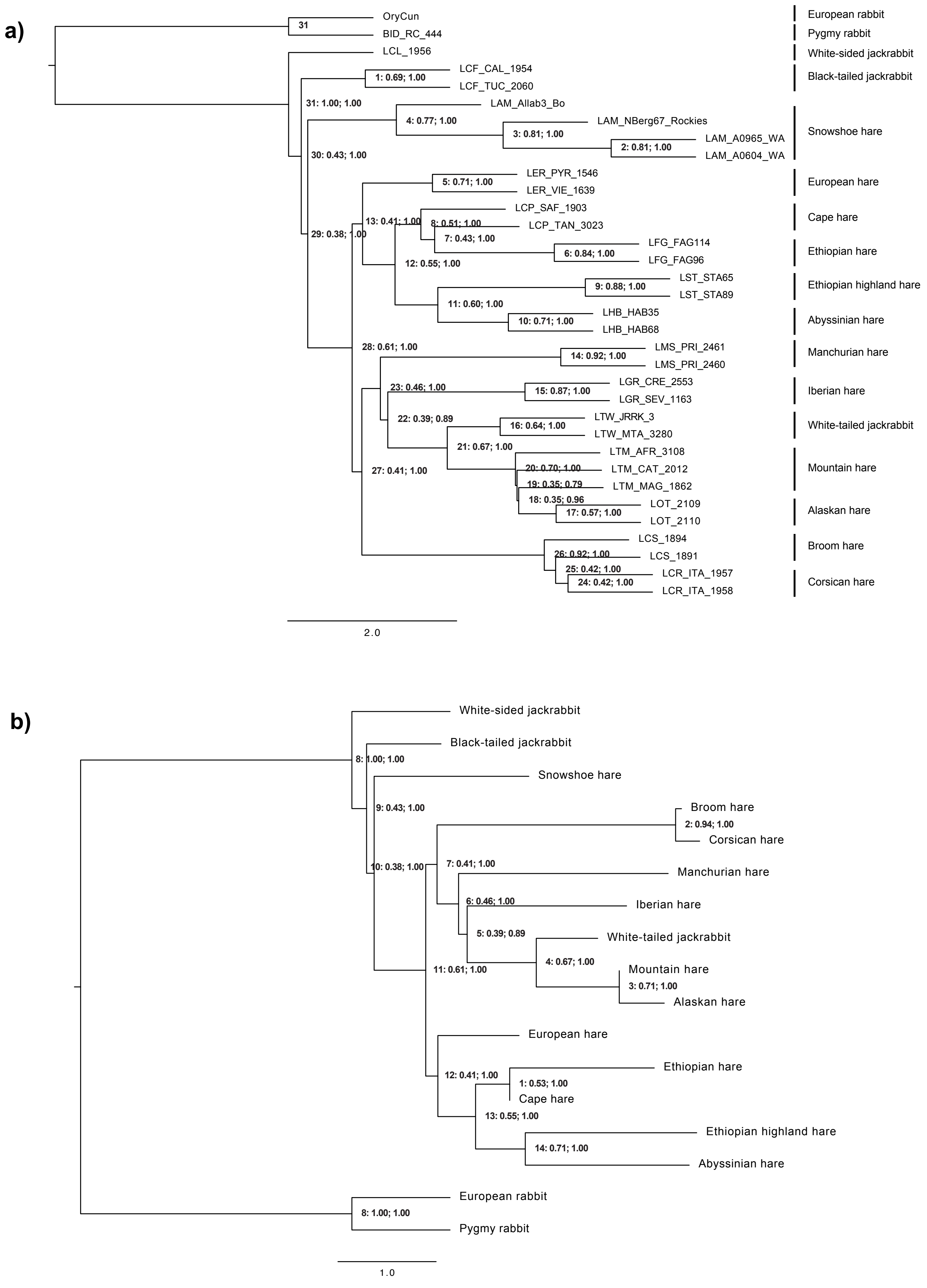


Figure S2 – *ASTRAL* species tree obtained with 8889 gene trees without a) or with b) assigning individuals to species. Nodes in both trees are numbered and annotated with quartet scores and posterior probabilities. Further information for each node can be found in Supplementary Tables S7 and S8.

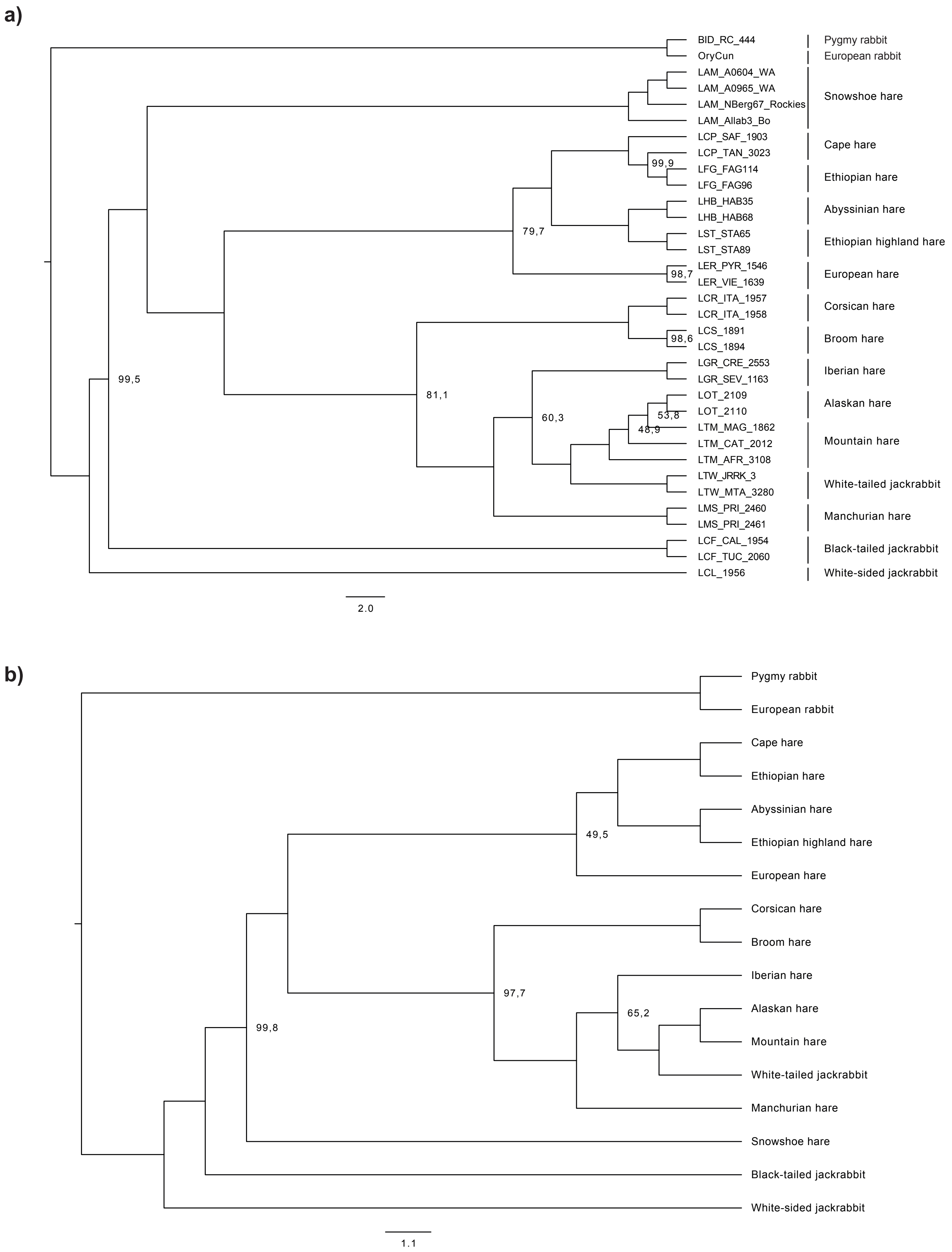


Figure S3 – *SVDquartets* species tree generated with 45,779 unlinked SNPs without a) or with b) assigning individuals to species. For both trees, we show branch supports for branches with support lower than 100% of 1000 bootstrap rounds.

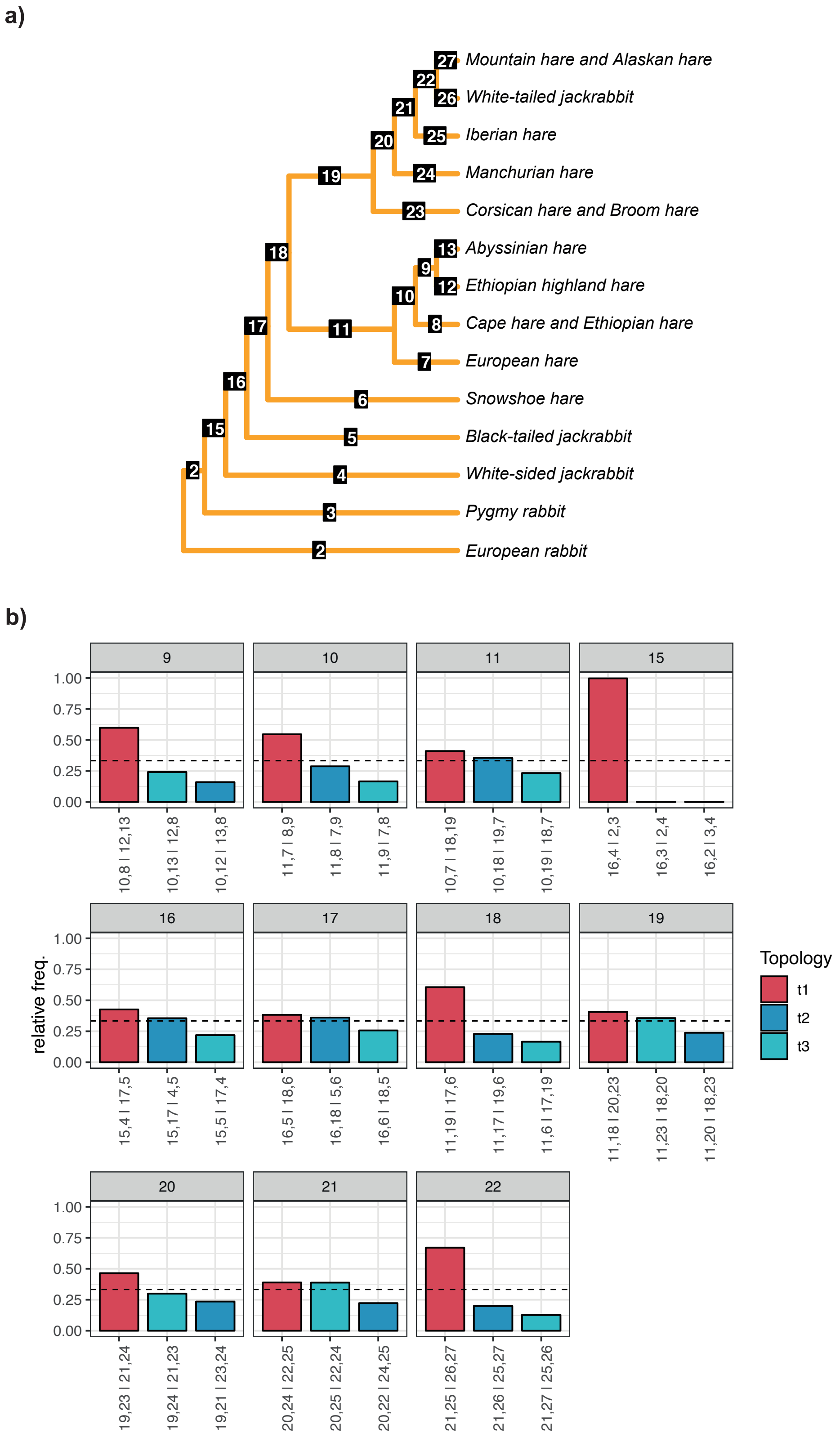


Figure S4 – Frequencies of the three main topologies around focal branches of the *ASTRAL* species tree (Fig. 1b). For each internal branch of the tree in (a), the frequency of the three possible topologies connecting the four neighboring branches is shown in (b). The title of each bar plot in (b) corresponds to labeled internal branches in the tree in (a). The most frequent topology is shown in red, and the two alternative topologies are shown in blue. On the x-axis of each bar plot, the topology of each quartet is represented using the neighboring branch labels. The branches leading to Mountain hares and Alaskan hares, Cape and Ethiopian hares, and Corsican and Broom hares were collapsed given the non-monophyly of the individuals of these species (Supplementary Figs. S2a and S3a).

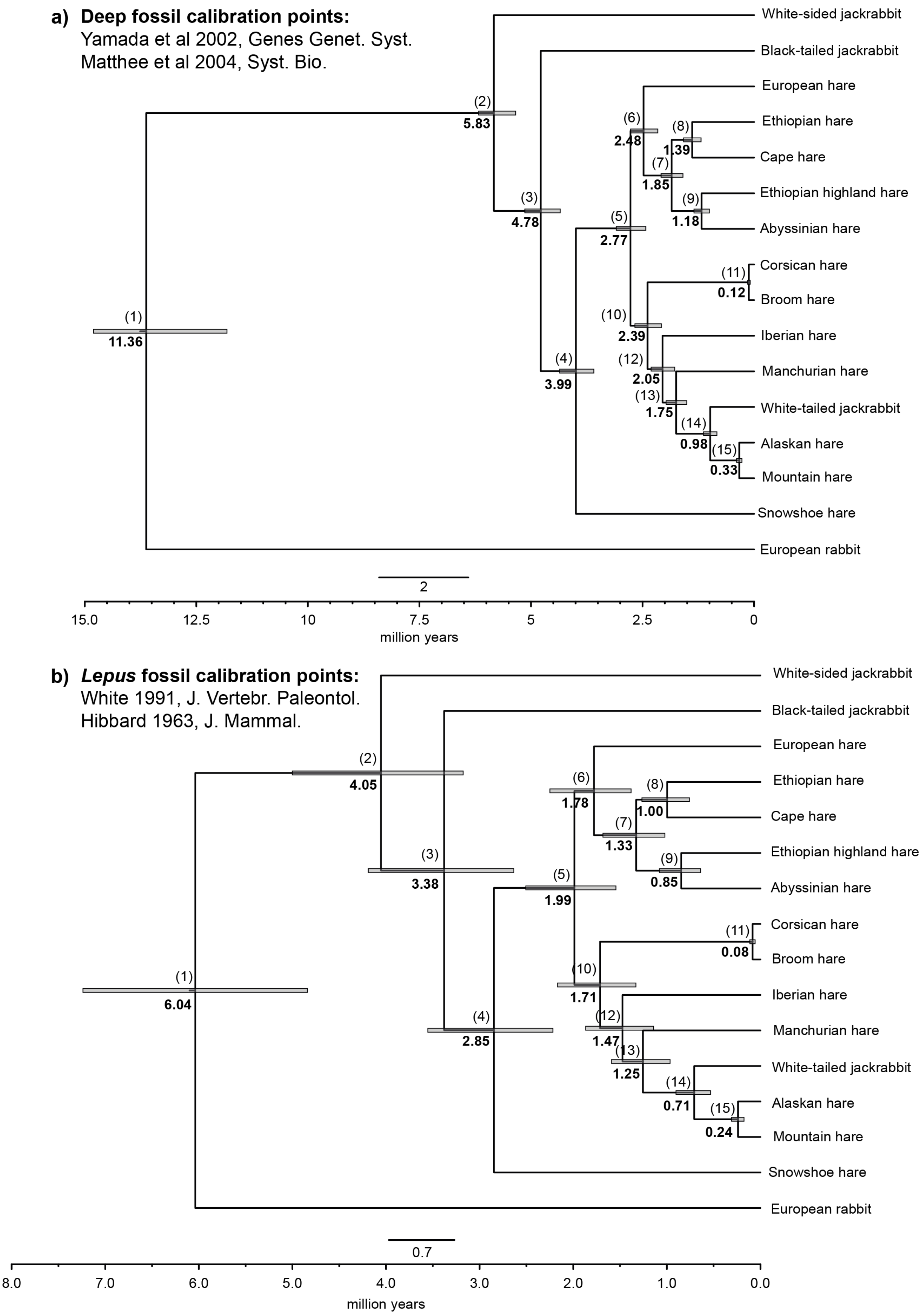


Figure S5 - Divergence time tree of *Lepus,* estimated from alignments of 9015 protein coding ortholog genes (10,863,822 bp alignment), with three partitions (1st, 2nd and 3rd codon positions) using (a) molecular-based dates extrapolated from deep fossil calibrations in the lagomorphs or (b) fossil calibrations for the divergence of the genus *Lepus* and the *Lepus* species (see Materials and Methods). Node labels in bold black are divergence time estimations in millions of years. Numbers in parenthesis in each node represent node numbers in Supplementary Table S5.

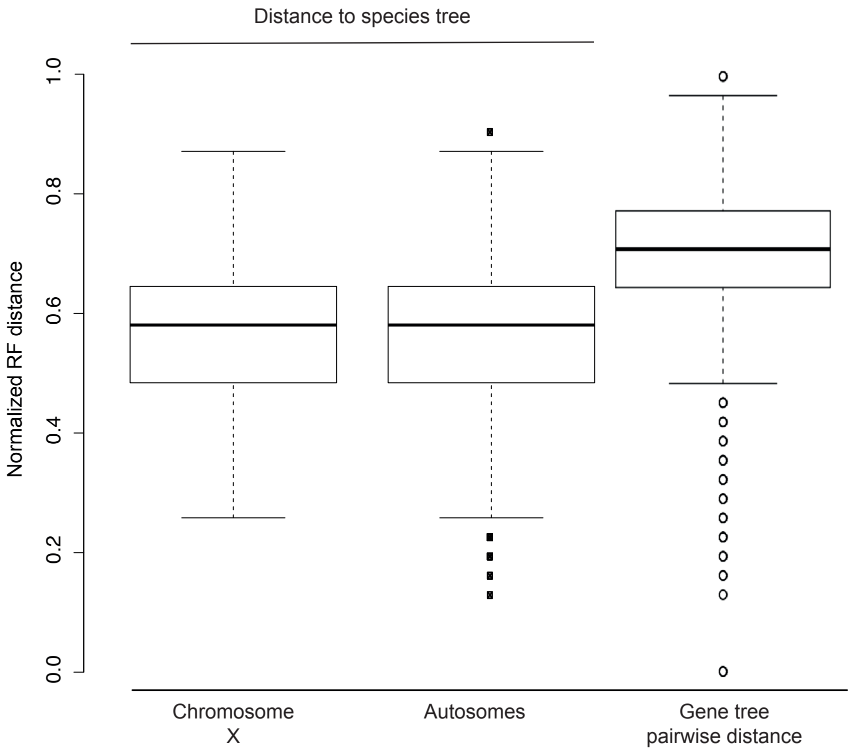


Figure S6 – Box plots of the normalized Robinson-Foulds distance between chromosome X and autosomes gene trees and the genome-wide species tree (Supplementary Fig. S2a), and pairwise distance among all gene trees.

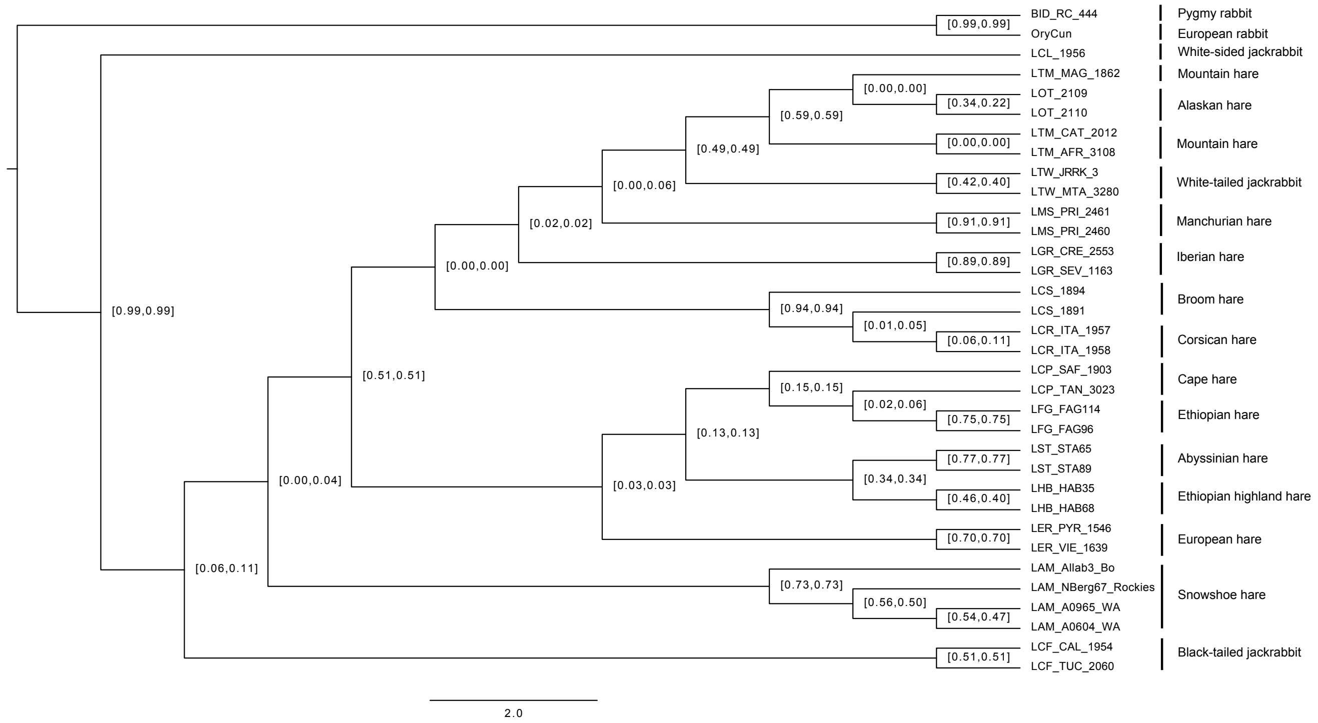


Figure S7 – Majority Rule Consensus Tree constructed from 8889 maximum likelihood gene trees. Tree nodes are annotated with the Internode Certainty (IC) and Internode Certainty All (ICA).

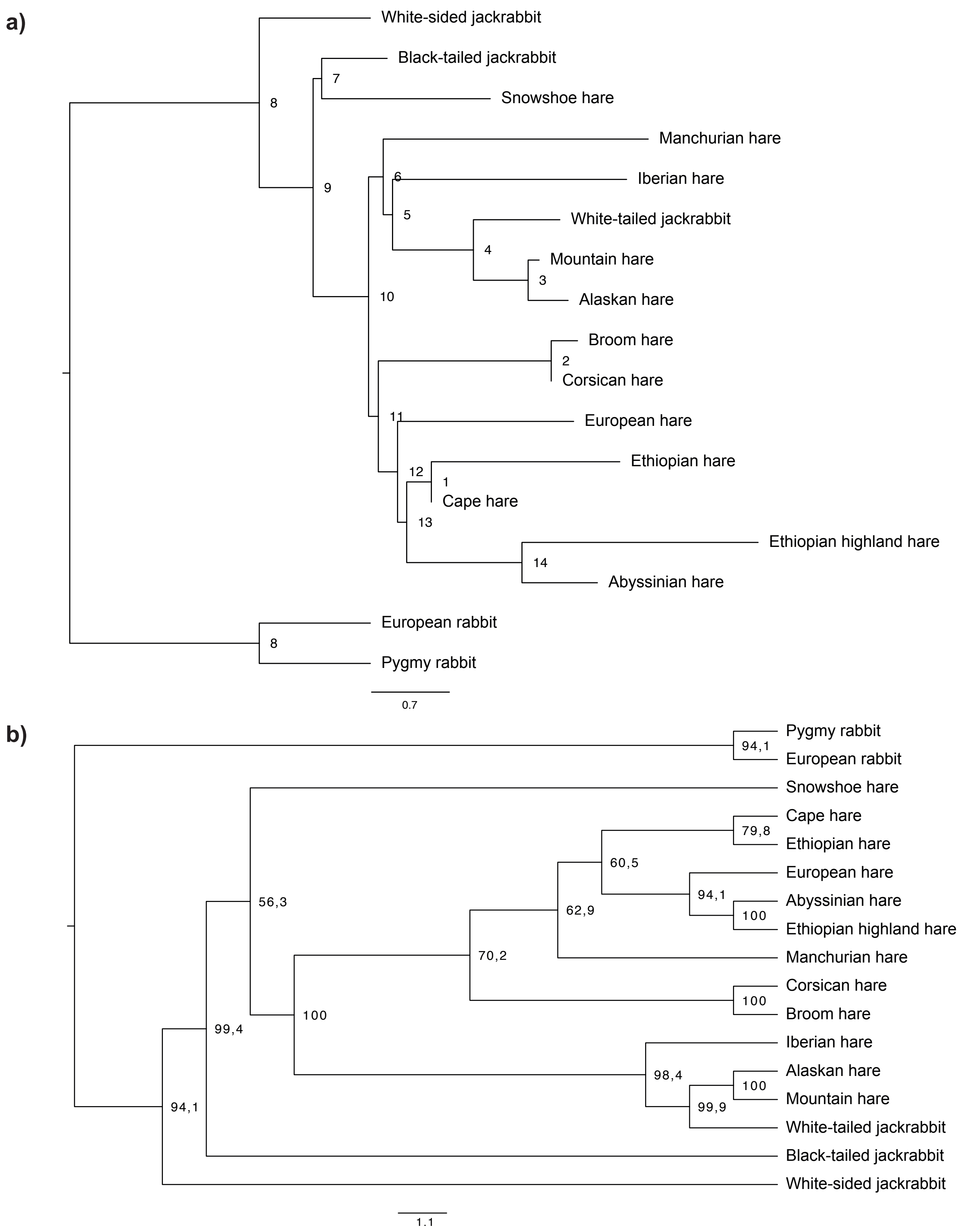


Figure S8 – Chromosome X species trees. (a) *ASTRAL* species tree obtained from 181 chromosome X gene trees and (b) *SVDquartets* species obtained from 1473 unlinked SNPs localized in the X chromosome. Node labels in (a) correspond to node labels in Supplementary Table S9 and (b) 1000 bootstrap support.

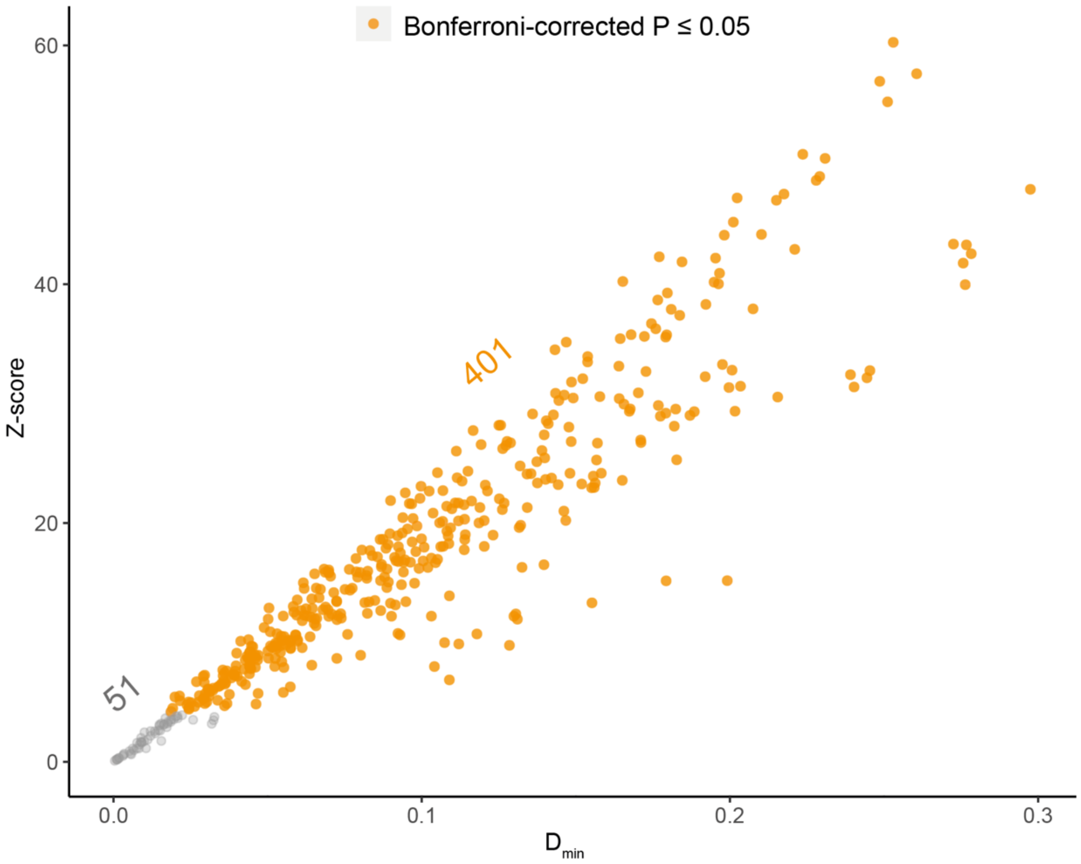

Figure S9 – Minimum D-statistic ($D_{min}$; Malinksy et al 2018, Nat. Ecol. Evol.) between all possible trios of taxa in the species tree plotted against their z-score. Significant values are highlighted in orange. The number of significant and non-significant $D_{min}$ values is given above the graph.

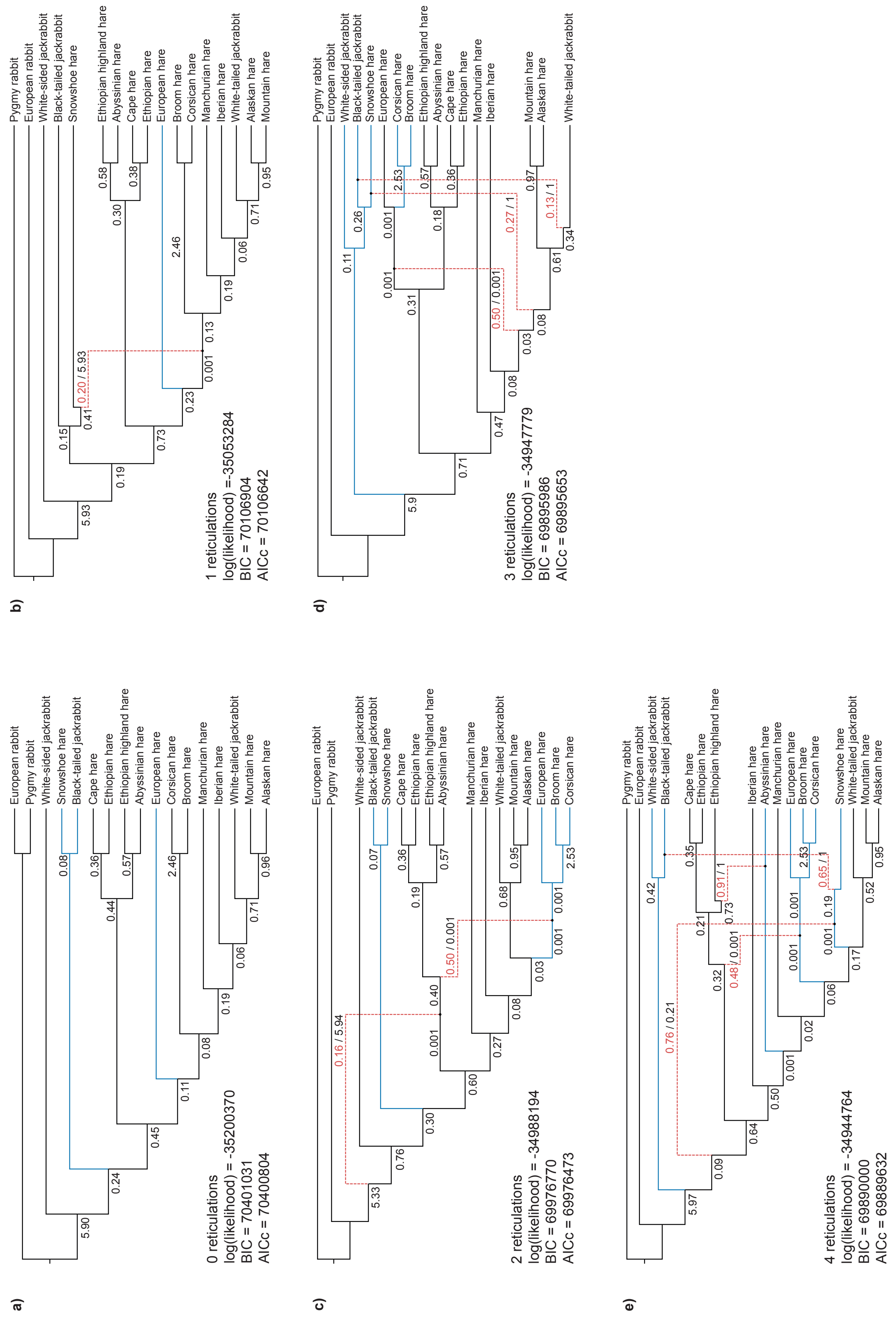


Figure S10 – Rooted pseudo-maximum likelihood networks for *Lepus*, with 0 to 4 reticulations inferred from 8889 gene trees with *PhyloNet*. Red branches represent reticulations, and blue branches represent variations to the species tree topology in Figure 1b. Branch lengths are represented in black and inheritance probabilities are represented in red. Bellow each network, we report the log likelihood, Bayesian information criteria (BIC) and Akaike information criteria corrected for small sample sizes (AICc).

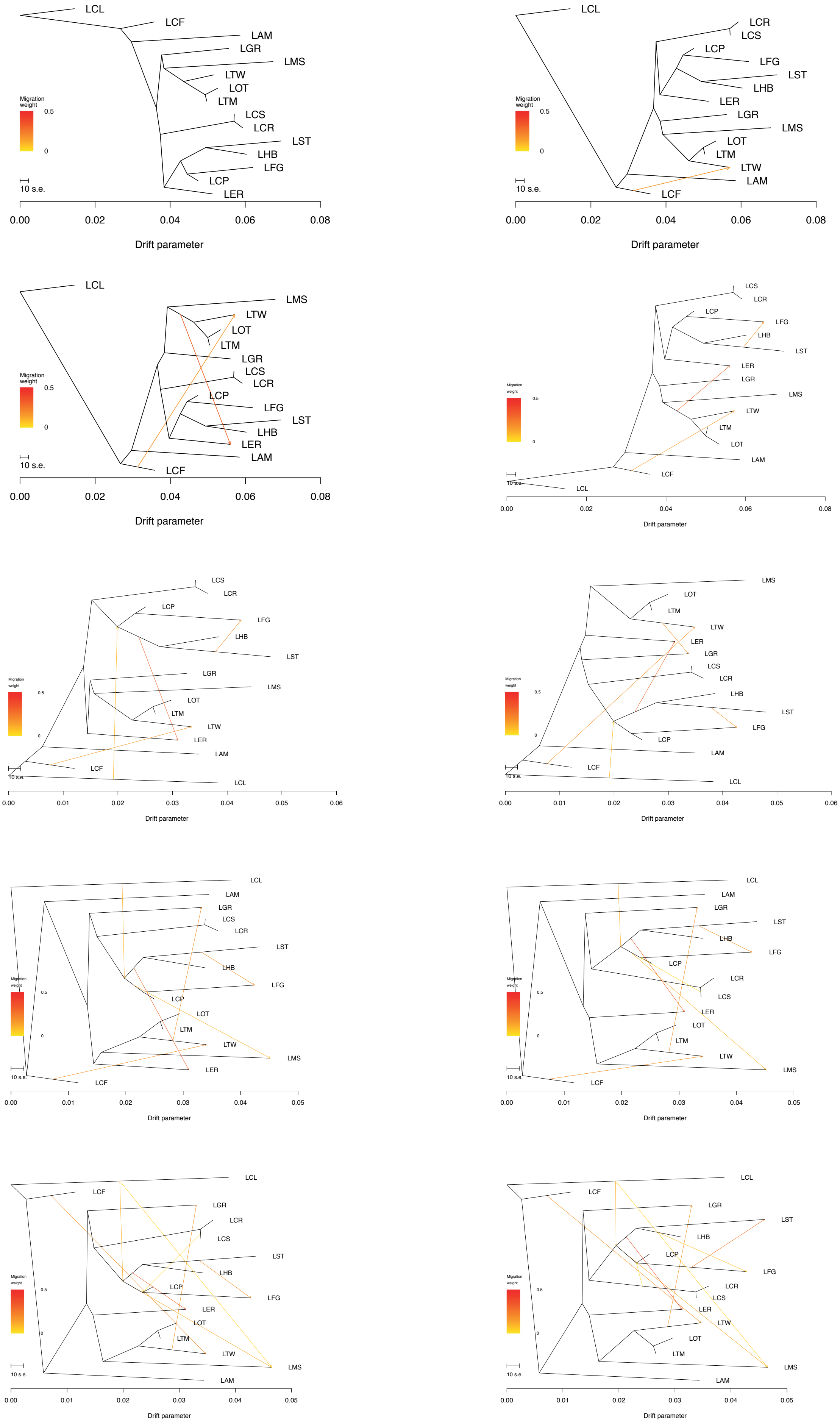
LCL
LCF
LAM
LGR
LMS
LTW
LOT
LTM
LCS
LCR
LST
LHB
LFG
LCP
LER
Migration weight
0.5
0
10 s.e.
0.00
0.02
0.04
0.06
0.08
Drift parameter
LCL
LCR
LCS
LCP
LFG
LST
LHB
LER
LGR
LMS
LOT
LTM
LTW
LAM
LCF
Migration weight
0.5
0
10 s.e.
0.00
0.02
0.04
0.06
0.08
Drift parameter
LCL
LMS
LTW
LOT
LTM
LGR
LCS
LCR
LCP
LFG
LST
LHB
LER
LAM
LCF
Migration weight
0.5
0
10 s.e.
0.00
0.02
0.04
0.06
0.08
Drift parameter
LCS
LCR
LCP
LFG
LHB
LST
LER
LGR
LMS
LTW
LTM
LOT
LAM
LCF
LCL
Migration weight
0.5
0
10 s.e.
0.00
0.02
0.04
0.06
0.08
Drift parameter
LCS
LCR
LCP
LFG
LHB
LST
LGR
LMS
LOT
LTM
LTW
LER
LAM
LCF
LCL
Migration weight
0.5
0
10 s.e.
0.00
0.01
0.02
0.03
0.04
0.05
0.06
Drift parameter
LMS
LOT
LTM
LTW
LER
LGR
LCS
LCR
LHB
LST
LFG
LCP
LAM
LCF
LCL
Migration weight
0.5
0
10 s.e.
0.00
0.01
0.02
0.03
0.04
0.05
0.06
Drift parameter
LCL
LAM
LGR
LCS
LCR
LST
LHB
LFG
LCP
LOT
LTM
LTW
LMS
LER
LCF
Migration weight
0.5
0
10 s.e.
0.00
0.01
0.02
0.03
0.04
0.05
Drift parameter
LCL
LAM
LGR
LST
LHB
LFG
LCP
LCR
LCS
LER
LOT
LTM
LTW
LMS
LCF
Migration weight
0.5
0
10 s.e.
0.00
0.01
0.02
0.03
0.04
0.05
Drift parameter
LCL
LCF
LGR
LCR
LCS
LST
LHB
LCP
LFG
LER
LOT
LTM
LTW
LMS
LAM
Migration weight
0.5
0
10 s.e.
0.00
0.01
0.02
0.03
0.04
0.05
Drift parameter
LCL
LCF
LGR
LST
LHB
LCP
LFG
LCR
LCS
LER
LTW
LOT
LTM
LMS
LAM
Migration weight
0.5
0
10 s.e.
0.00
0.01
0.02
0.03
0.04
0.05
Drift parameter

Figure S11

Figure S11 (continued) – *TreeMix* inference of splits and admixture using 30,709 biallelic SNPs for no migration events (m=0) to nine migration events (m=9). The white-sided jackrabbit (LCL) was used as outgroup. LAM – Snowshoe hare; LCF – Black-tailed jackrabbit; LGR – Iberian hare; LCR – Corsican hare; LCS – Broom hare; LST – Ethiopian highland hare; LHB – Abyssinian hare; LCP – Cape hare; LFG – Ethiopian hare; LER – European hare; LOT – Alaskan hare; LTM – Mountain hare; LTW – White-tailed jackrabbit; LMS – Manchurian hare.

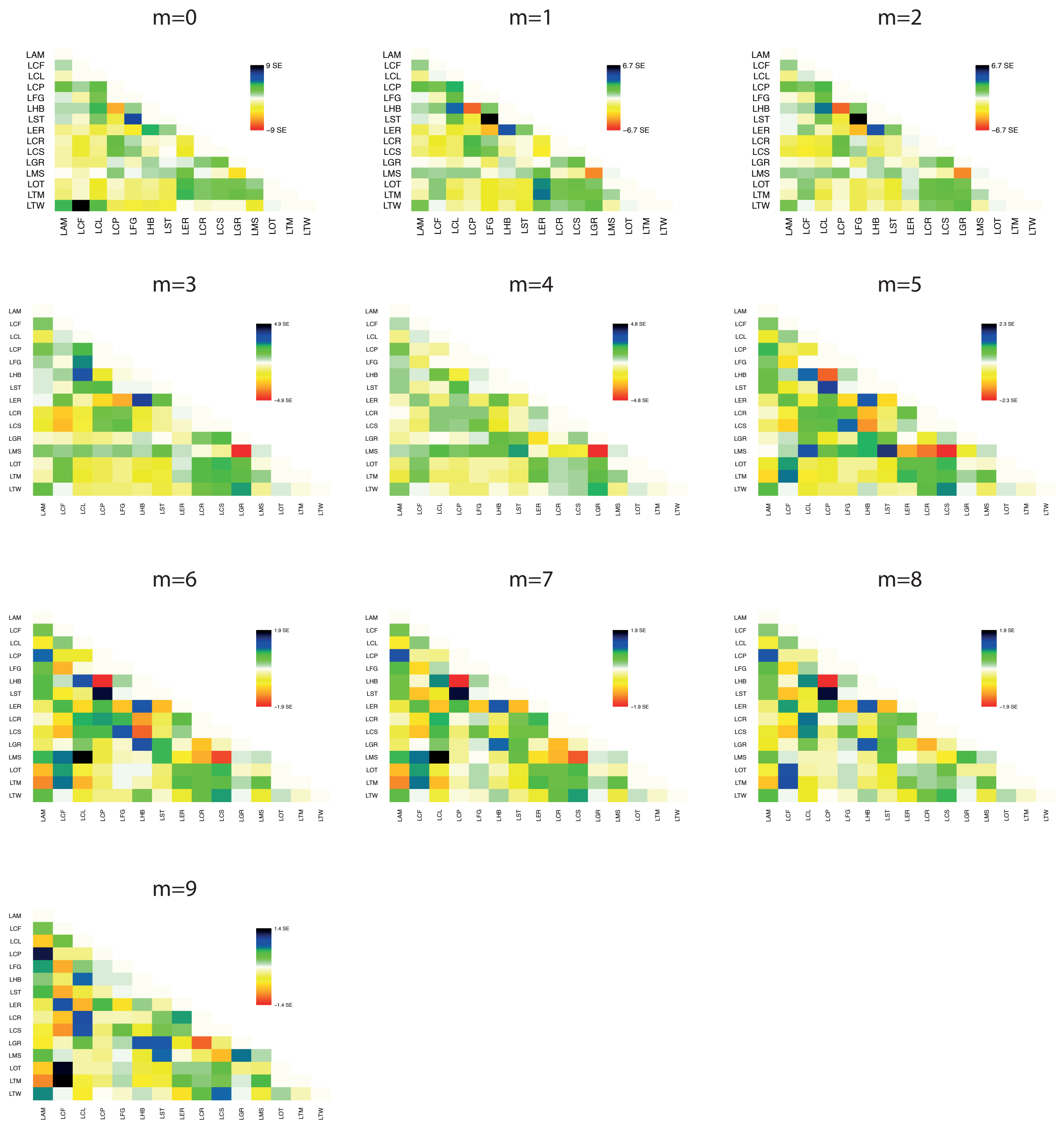


Figure S12 – Residuals for the *TreeMix* networks of Supplementary Figure S11, from 0 to 9 migration events. The white-sided jackrabbit (LCL) was used as outgroup. LAM – Snowshoe hare; LCF – Black-tailed jackrabbit; LGR – Iberian hare; LCR – Corsican hare; LCS – Broom hare; LST – Ethiopian highland hare; LHB – Abyssinian hare; LCP – Cape hare; LFG – Ethiopian hare; LER – European hare; LOT – Alaskan hare; LTM – Mountain hare; LTW – White-tailed jackrabbit; LMS – Manchurian hare.

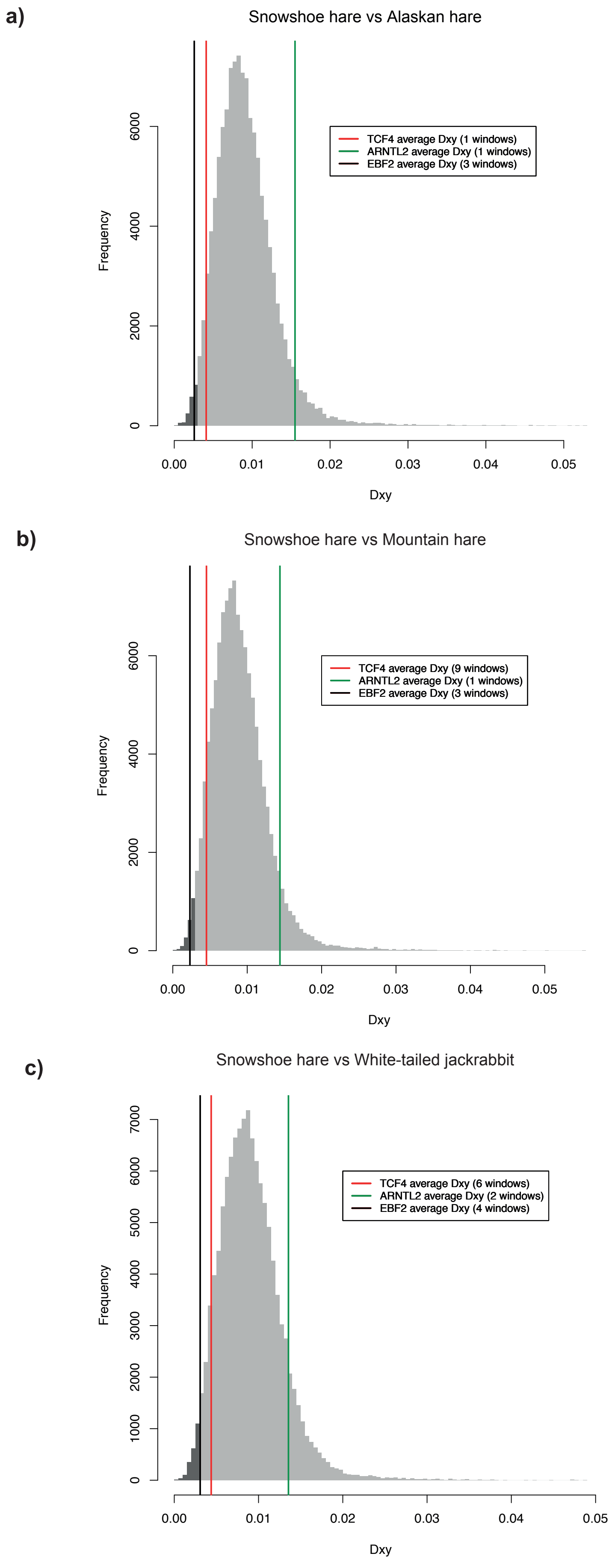


Figure S13 – Exome-wide observed genetic divergence (gray distribution) and average genetic divergence (colored lines) for fraction of admixture ($f_d$) outlier windows (Supplementary Table S12) between snowshoe hares and (a) Alaskan hare, (b) Mountain hare and (c) white-tailed jackrabbits. The number of outlier windows used to estimate $d_{xy}$ is reported above the graph, and in dark gray we highlight the lower 1% percentile of the distribution.
